# The evolutionary history of the common bean (*Phaseolus vulgaris*) revealed by chloroplast and nuclear genomes

**DOI:** 10.1101/2023.06.09.544374

**Authors:** Giulia Frascarelli, Teresa R. Galise, Nunzio D’Agostino, Donata Cafasso, Salvatore Cozzolino, Gaia Cortinovis, Francesca Sparvoli, Elisa Bellucci, Valerio Di Vittori, Laura Nanni, Alice Pieri, Marzia Rossato, Leonardo Vincenzi, Andrea Benazzo, Massimo Delledonne, Elena Bitocchi, Roberto Papa

## Abstract

The remarkable evolutionary history of the common bean (*Phaseolus vulgaris L.*) has led to the emergence of three wild main genepools corresponding to three different ecogeographic areas: Mesoamerica, the Andes and northern Peru/Ecuador. Recent works proposed novel scenarios and the northern Peru/Ecuador population has been described as a new species called *P. debouckii,* rekindling the debate about the origin of *P. vulgaris*. Here we shed light on the origin of *P. vulgaris* by analysing the chloroplast and nuclear genomes of a large varietal collection representing the entire geographical distribution of wild forms. We assembled 37 chloroplast genomes *de novo* and used them to construct a time frame for the divergence of the genotypes under investigation, revealing that the separation of the Mesoamerican and northern Peru/Ecuador genepools occurred ∼0.15 Mya. Our results clearly support a Mesoamerican origin of the common bean and reject the recent *P. deboukii* hypothesis. These results also imply two independent migratory events from Mesoamerica to the North and South Andes, probably facilitated by birds. Our work represents a paradigmatic example of the importance of taking into account recombination events when investigating phylogeny and of the analysis of wild forms when studying the evolutionary history of a crop species.

## Introduction

Wild forms of the common bean (*Phaseolus vulgaris L.*) grow over a large geographic area in Latin America, from northern Mexico to north-western Argentina (Toro et al. 1990). The species has an intriguing evolutionary history that has resulted in at least three eco-geographical genepools: Mesoamerican, Andean and northern Peru/Ecuador. The Mesoamerican genepool is distributed in Mexico, Central America, Colombia and Venezuela, while the Andean genotypes spread from southern Peru to Bolivia and Argentina. Conversely, the northern Peru/Ecuador genepool identified by Debouck (1986) is restricted to the western Andes. The first two populations include both wild and domesticated forms, whereas the northern Peru/Ecuador genepool only has wild forms. Phylogeographic analysis, which emphasizes the close connection between genealogy and geography (Avise at al., 1986), has shown that the domestication of the common bean was preceded by two geographically distinct and isolated lineages (Mesoamerican and Andean). The genetic variability characteristics of wild relatives represents the primary resource for phylogenetics investigation. Indeed, domesticated genotypes underwent a genetic bottleneck due to the domestication process that resulted in a reduction of genetic variation and dramatic phenotypic changes (Meyer et al., 2012)

Although phylogenetics is necessary to infer evolutionary relationships (Cavalli-Sforza and Edwards, 1966), the rearrangement of genetic material during recombination events can produce artefacts (Schierup and Hein, 2000). Indeed, recombination implies that different parts of a sequence can have distinct phylogenetic histories and can be related by more than one tree (Nordborg and Tavaré, 2002). Furthermore, when considering recent divergence between species or populations, incomplete lineage sorting can cause further difficulties in the phylogenetic reconstruction and discordance between gene trees but also between nuclear and plastid trees. To address this issue, mitochondrial and chloroplast genomes have been extensively used to reconstruct the genetic lineages of animals and plants, respectively. Both organelles are haploid, implying a smaller population size than the nuclear genome, so organelle and nuclear genome data may trace different evolutionary histories. In plant genealogy, the chloroplast genome is widely preferred because it evolves more slowly than animal mitochondrial DNA but faster than plant mitochondrial DNA (Avise, 2009).

The origin of the common bean is still debated, and numerous theories have been advanced. The northern Peru/Ecuador hypothesis suggested by Kami et al. (1995) was based on the type I seed storage protein phaseolin, which is characteristic of a central area in northern Peru/Ecuador. The lack of tandem repeats in this type of phaseolin implied its ancestral state and supported northern Peru/Ecuador as the centre of origin. Evidence for a Mesoamerican origin was first provided by Rossi et al. (2009), who analysed the genetic diversity of the Mesoamerican and Andean gene pools using amplified fragment length polymorphism markers. Bitocchi et al. (2012) provided further support by investigating the origin of the three wild gene pools using nuclear single nucleotide polymorphisms (SNPs) at five independent loci, and their work was confirmed by the analysis of chloroplast simple sequence repeats (Desiderio et al 2013). More recently, a speciation event occurring before the divergence of the Mesoamerican and Andean genepools was hypothesized by Rendón-Anaya et al. (2017). The speciation gave rise to the northern Peru/Ecuador population that has been described by the authors as a new species of *Phaseolus* named *Phaseolus deboukii* (Rendón-Anaya et al., 2017). Based on the proposed Mesoamerican origin, Ariani et al. (2018) suggested the existence of a common ancestral population for the three genepools that became extinct when the Mesoamerican and Andean gene pools diverged (“Protovulgaris hypothesis”). We therefore set out to elucidate the evolutionary history *of P. vulgaris* by investigating the relationships between the three major wild genepools: Mesoamerican, Andean and northern Peru/Ecuador. We used chloroplast and nuclear sequence data of wild accessions, belonging to the three genepools, to reconstruct the phylogeny of this species and to infer times of divergence among the three wild genepools.

## Results

### Identification and analysis of SNP variation in the chloroplast genome

We selected 70 *P. vulgaris* wild accessions covering the geographical distribution of wild common bean from northern Mexico to north-western Argentina, thus representing all three genepools (44 Mesoamerican, 22 Andean and 4 northern Peru/Ecuador, the latter with the ancestral type I phaseolin protein). We also selected wild samples of *P. coccineus* (22) and *P. lunatus* (3), as well as one wild and one domesticated accession of *P. acutifolius*. Across the 97 *Phaseolus* samples, we identified 4008 SNPs (including 777 singletons), 1999 of which were found within genes. Among the 128 chloroplast genes, 66% (84) contained at least one SNP and 45% (56) contained more than three. The most variable genes were ndhF, accD and the pseudogene ycf1b, with 100, 115 and 391 SNPs, respectively. We found 1526 SNPs within exons and 473 within introns, and in the psbH gene all variants were located only within exons. Most of the SNPs were distributed across the single copy regions. A higher SNP density was found in the small single copy region (522 SNPs per 100 bp window on average) compared to the large single copy region (376 SNPs per 100 bp window on average). We found that 42.3% of the variants were synonymous and 57.7% were non-synonymous, with 56.97% of the non-synonymous variants resulting in missense mutations.

### Chloroplast genetic structure

Characteristics of chloroplast DNA such as uniparental inheritance, haploidy and lack of recombination make it more suitable than nuclear DNA for the reconstruction of intraspecific phylogenetic relationships. We analysed the chloroplast diversity of a large sample of ∼100 wild *Phaseolus* accessions, specifically n = 70 *P. vulgaris*, n = 22 *P. coccineus*, n = 2 *P. acutifolius*, and n = 3 *P. lunatus* (Supplementary Table 1). We also prepared de novo assemblies of 37 aligned chloroplast genomes (Supplementary Figures 1–4) and used them to corroborate the results obtained from artificial sequences prepared by the concatenation of SNPs. We used 3231 SNPs to investigate the genetic structure of the *P. vulgaris* chloroplast (Figure 1a,b). The whole population can be divided into five subgroups named S1–S5 (Figure 1b) based on a log marginal likelihood of optimal partition of – 6689.6434. All PhI samples in the panel were pooled in S1. The Mesoamerican accessions were split across S2, S3 and S5. Cluster S2 included three samples from Guatemala, Costa Rica and Honduras, whereas cluster S3 only included Mexican accessions and cluster S5 contained 25 samples from Mexico, two from Colombia and one from Guatemala. All Andean accessions were clustered in S4. Relationships between and within species were inspected by multidimensional scaling (MDS) based on an identity-by-state matrix used as a distance matrix. The MDS plot separated the 97 accessions by species (Supplementary Figure 5). Specifically, the first component (C1) separated *P. coccineus*, *P. acutifolius* and *P. lunatus*, whereas the second (C2) separated *P. vulgaris* and *P. coccineus*. Within the *P. vulgaris* group (Figure 1c), C1 separated the Andean and Mesoamerican genepools, and also divided the accessions belonging to the latter into three groups, one of which was closest to the Andean samples. C2 separated the PhI genepool from the Andean and Mesoamerican genepools and also separated accessions 59_Pv_MW_CR, 787a_Pv_MW_GT and 790_MW_HN from the rest of the Mesoamerican samples.

**Figure 1:**
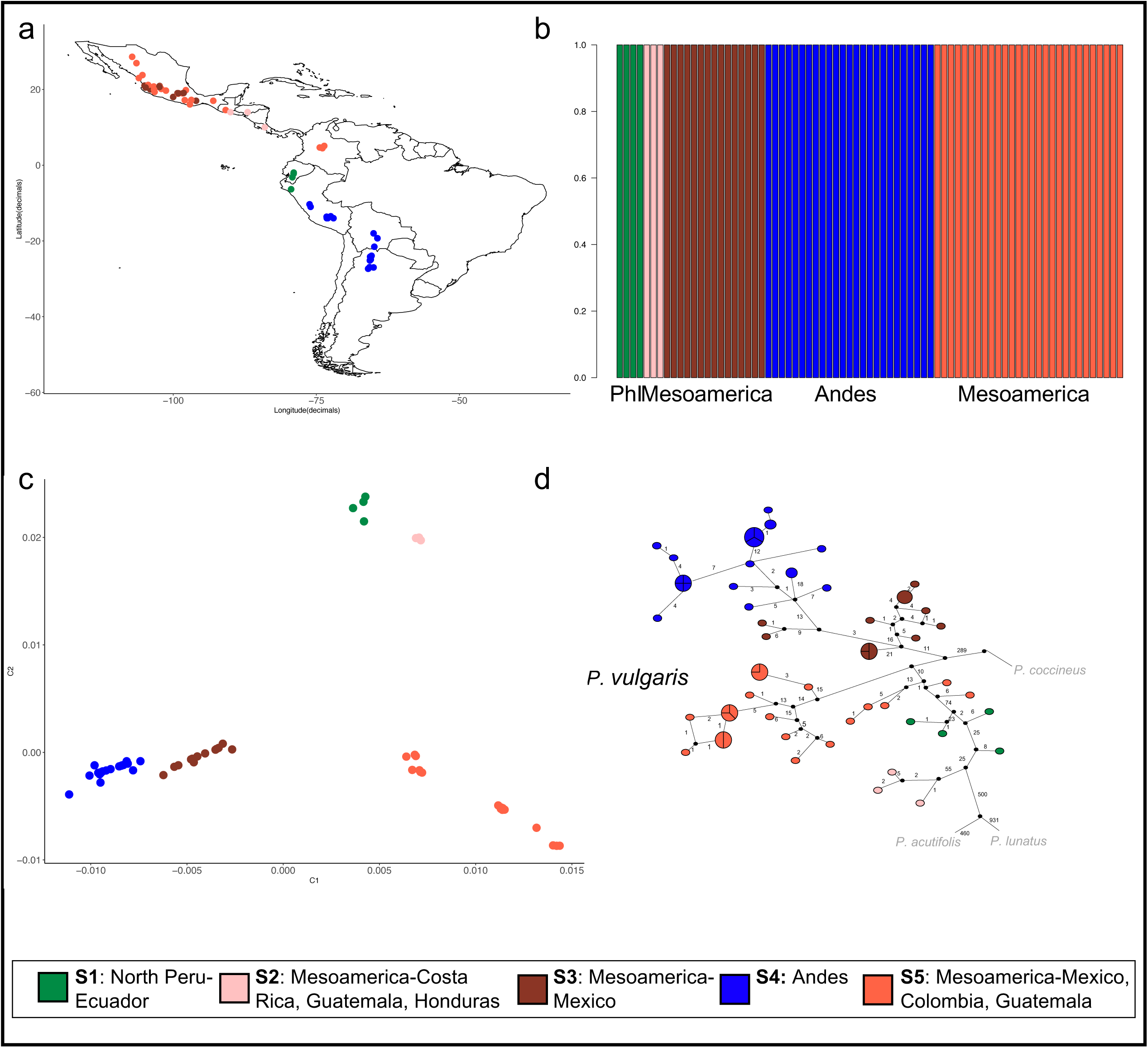
Analysis of the *P. vulgaris* population structure. (a) Geographical distribution of *P. vulgaris* accessions based on BAPS cluster membership. (b) Population structure. (c) MDS plot of *P. vulgaris* samples. (d) Haplotype network analysis of all *Phaseolus* accessions, focusing on *P. vulgaris*. MW – Mesoamerican wild; AW – Andean wild; PhI – Phaseolin type I. Each circle represents a single haplotype and the size of the circles is proportional to the number of individuals carrying the same haplotype. Black dots indicate missing intermediate haplotypes and numbers correspond to mutational steps.

Haplotype network analysis was used to visualize pedigree relationships at the intraspecific level. Forty-five haplotypes were identified within the species *P. vulgaris* (Figure 1d), and no haplotypes were shared between the Mesoamerican and Andean genepools. Within the Mesoamerican genepool, four Mexican and three Columbian accessions shared the same haplotype, as did two samples from Mexico and one from Guatemala. Furthermore, in the Andean genepool, two samples from Peru shared the haplotype with an accession from Bolivia and seven samples from Argentina. The accessions carrying phaseolin type I showed haplotypes that were mostly separated from the Andean samples and closest to the Mesoamerican genepool, particularly to the three accessions from Guatemala, Honduras and Costa Rica (787a_Pv_MW_GT, 790_Pv_MW_HN and 059_Pv_MW_CR, respectively).

### Phylogenetic analysis of chloroplast DNA

We investigated the phylogenetic relationships among the *P. vulgaris* accessions based on the alignment of the 37 *de novo* chloroplast genomes (Figure 2). The maximum-likelihood (ML) tree clearly showed the presence of a genetic structure in Mesoamerica with three well supported groups (bootstrap values > 70%). The first included accessions from Guatemala, Honduras and Costa Rica, and was more closely related to the northern Peru/Ecuador genepool. The second included only Mexican accessions pooled with Andean samples, and the third consisted of samples from Mexico, Guatemala and Colombia. The clade of accessions from northern Peru/Ecuador clearly arose from the Mesoamerican genepool. The ML tree constructed from SNP data (Supplementary Figure 6) was topologically consistent with the tree constructed from the chloroplast genomes.

**Figure 2:**
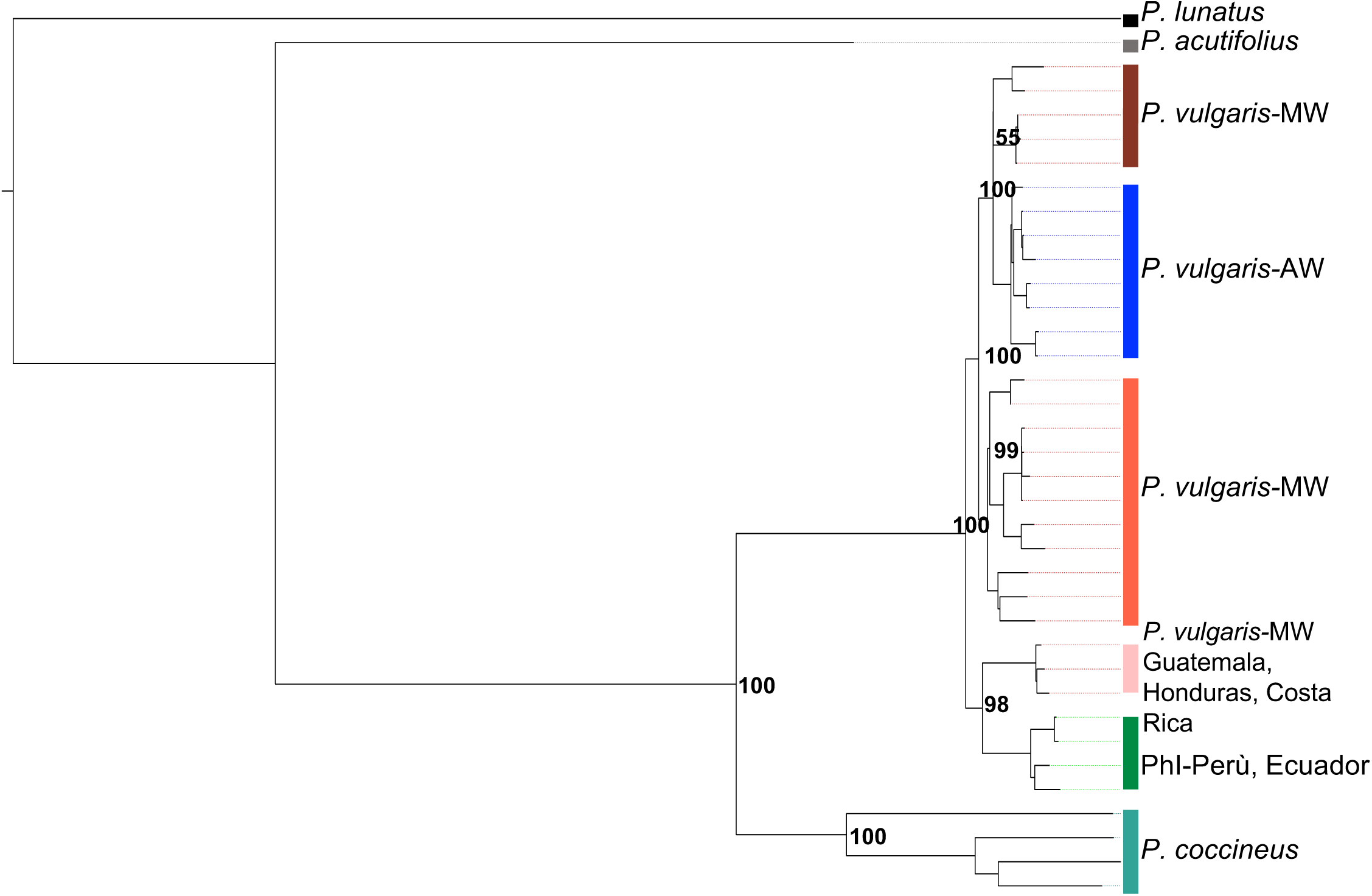
Maximum-likelihood tree obtained from the alignment of 37 chloroplast genomes (bootstrap value = 10,000). MW – Mesoamerican wild; AW – Andean wild.

### Molecular clock analysis of chloroplast DNA

To estimate timelines for the divergence of the *P. vulgaris* genepools, we combined the 37 *de novo* chloroplast genome assemblies with three *Vigna spp.* plastomes and carried out a coalescent simulation with a relaxed log-normal molecular clock, using the divergence between *P. coccineus* and *Vigna spp.* to calibrate the tree (Lavin et al., 2005). The coalescent simulation (Figure 3) showed a divergence time of ∼0.19 million years ago (Mya) between the wild *P. vulgaris* genetic groups (0.0847– 0.3082 95% highest posterior probability, HPD). The separation between the northern Peru/Ecuador genepool and the group comprising the Mesoamerican and Andean genepools occurred ∼0.15 Mya (0.0607–0.2419 95% HPD). A split between the Mesoamerican and Andean populations was very recent, at ∼0.09 Mya (0.0422-0.1515 95% HPD).

**Figure 3:**
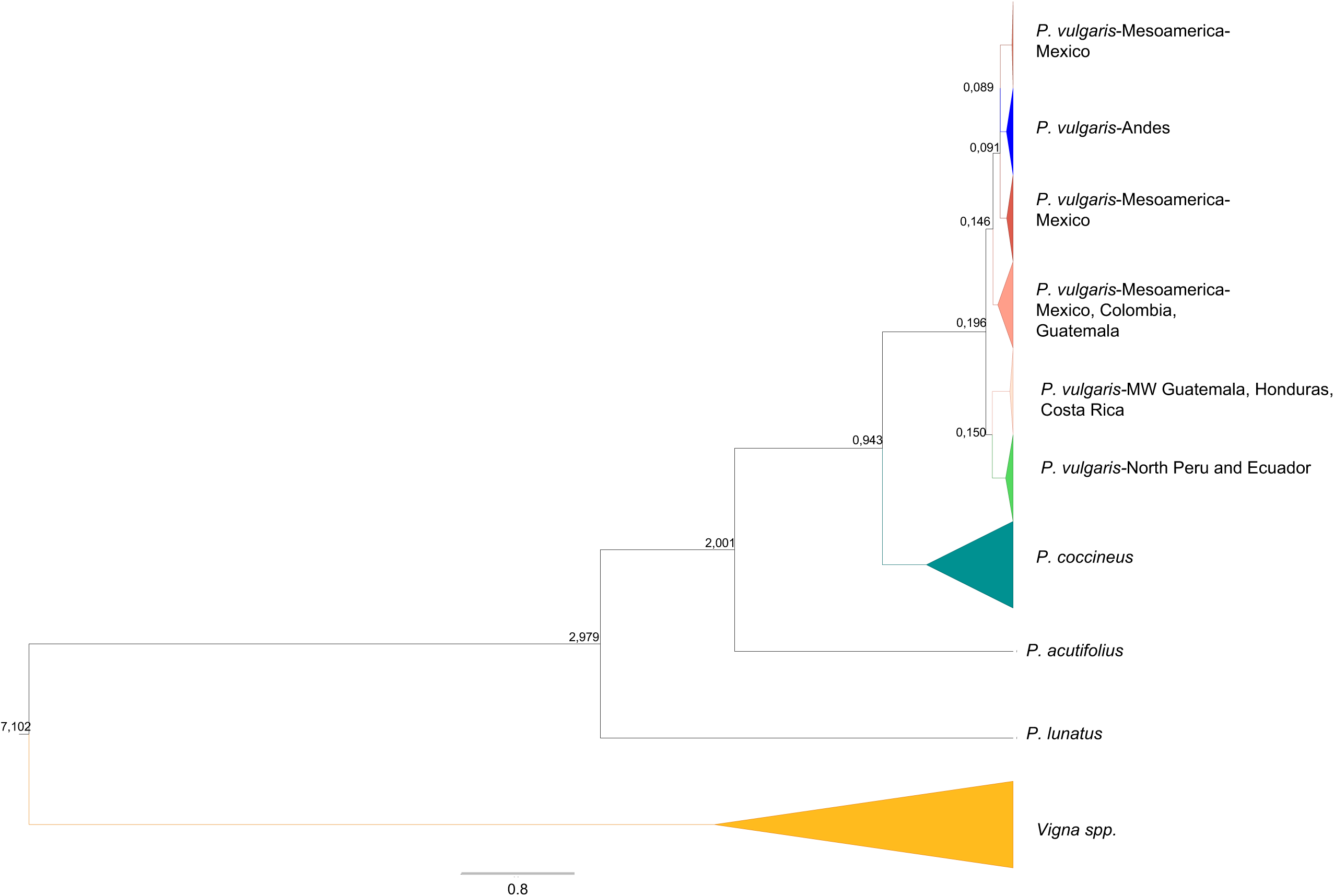
Molecular clock analysis computed using 37 chloroplast genomes. Divergence times are shown on the nodes. MW – Mesoamerican wild; AW – Andean wild; PhI – Phaseolin type I (northern Peru/Ecuador).

### Intraspecies phylogenetic analysis of the nuclear genome

Nuclear data from 10 *P. vulgaris* accessions representing the three wild genepools (Supplementary Table 2) were also analysed to infer phylogenetic relationships. We identified 11,160,422 SNPs, most of which were found in intergenic (45.30%), upstream (19.88%) or downstream (20.00%) regions. A further 7.45% of the SNPs were found within introns and 4.86% within exons. Only SNPs located in the pericentromeric regions, which are less prone to recombination, were used to construct an ML tree for each of the 11 chromosomes (Supplementary Table 3).

Although different topologies were obtained for the 11 chromosome-specific ML trees (Figure 4), most nodes were well supported (bootstrap values > 70%). No clear distinction was observed between the Mesoamerican and Andean genepools. Interestingly, PhI and Mesoamerican samples from Costa Rica, Guatemala and Mexico (Oaxaca) grouped together (078_Pv_PhI_EC, 059_Pv_MW_CR, 787a_Pv_MW_GT and 081_Pv_MW_MX). We also constructed a neighbour-joining tree from concatemers of genome-wide SNPs pruned every 250 kb (Supplementary Figure 7). Contrary to the results obtained with chloroplast sequences, the nuclear data were insufficient to make inferences about the origin of and divergence between the three wild genepools.

**Figure 4:**
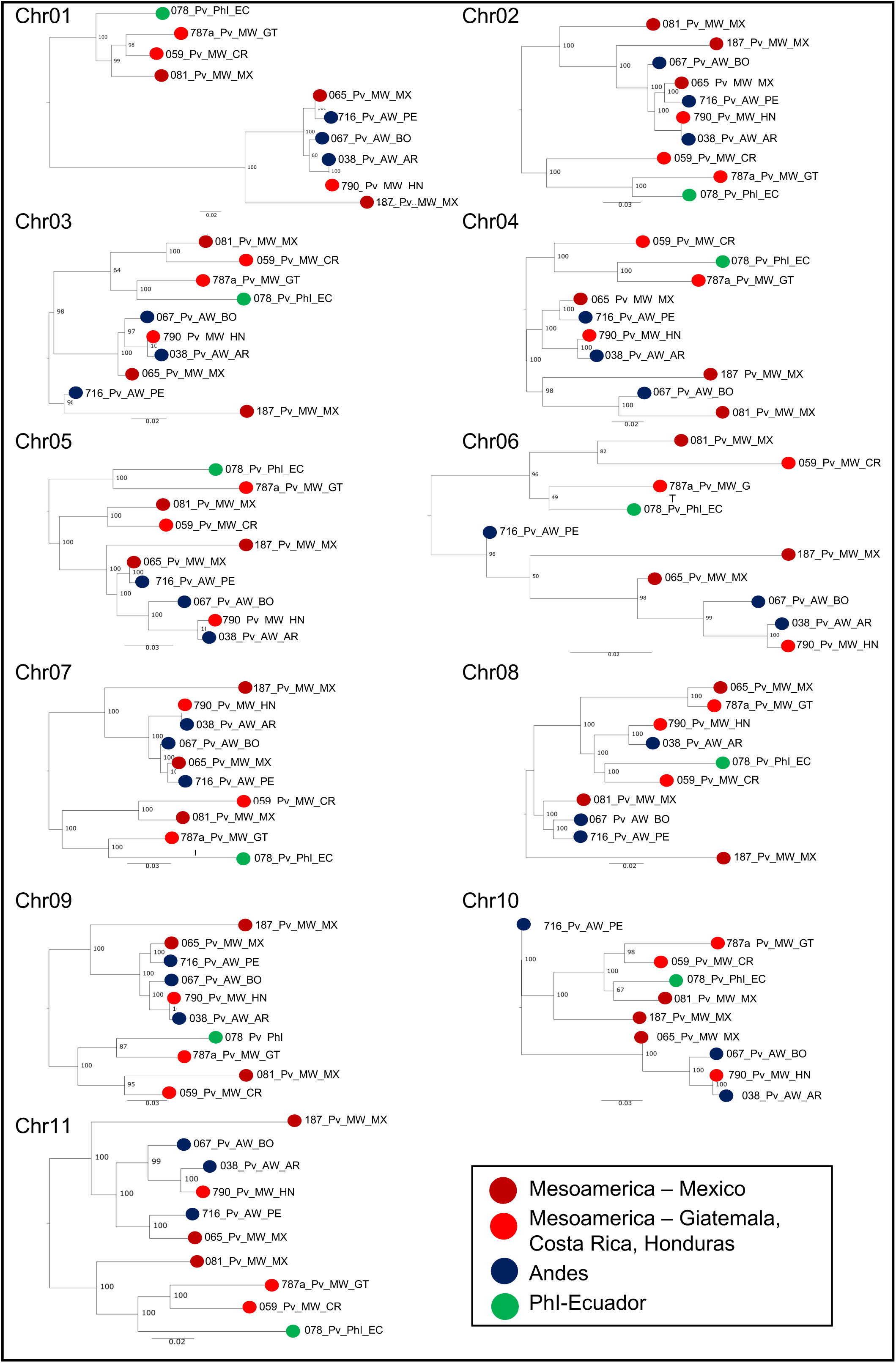
Maximum-likelihood trees based on concatenated SNPs extracted from the centromeric regions of the 11 chromosomes. Dark red: Mesoamerican samples from Mexico. Light red: Mesoamerican samples from Guatemala, Costa Rica or Honduras. Blue: Andean samples. Green: PhI sample from Ecuador.

## Discussion

Knowledge of the origin, evolution and diffusion of crops is necessary for the appropriate use and conservation of available genetic resources. Wild forms are characterized by extensive genetic and phenotypic diversity, which firstly must be recovered and exploited to accelerate breeding programs (Gepts, 1990) and secondly, its study allows to trace back the evolutionary history of a species.

Indeed, reconstructing the evolutionary events that occurred before domestication gets difficult if only domesticated forms/genotypes are considered, due to gene flow between wild and domesticated forms (Kwak et al., 2009) admixture between different cultigens and the reduction of the genetic variability present in the crops, caused by the domestication process itself and becomes hard to reconstruct a reliable population structure of the wild forms.

In this study, we carried out an intraspecific phylogenetic analysis based on chloroplast and nuclear sequence data to elucidate the relationships between wild *Phaseolus vulgaris* genepools. We analysed the genetic diversity of a large collection of *P. vulgaris* chloroplast genomes representing the three wild genepools. A selection of these genotypes was then used to identify nuclear SNPs suitable to reconstruct the phylogeography of the common bean and clarify the relationships and timeframe for the divergence of the wild genetic groups.

Our results assigned PhI accessions from northern Peru/Ecuador to a clade that clearly arose from the Mesoamerican genepool, in agreement with earlier reports (Bitocchi et al., 2012). This was supported by complementary phylogenesis, Bayesian analysis of population structure (BAPS), MDS, and haplotype network analysis, revealing a clear subdivision of the Mesoamerican population. Two of the Mesoamerican subpopulations were placed closest to the Andean genepool and PhI accessions, respectively. Similar results were reported by Bitocchi et al. (2012) and Desiderio et al. (2013) using different molecular markers. Our findings therefore support the monophyletic and Mesoamerican origin of the common bean, and revealed no evidence supporting PhI population speciation, as proposed by Rendón-Anaya (2017). The estimated divergence times between *P. vulgaris* genepools provided additional supporting evidence. The coalescent simulation showed a divergence time between wild *P. vulgaris* groups of ∼0.19 Mya.

Our estimate of ∼0.19 Mya overlaps with that computed by Rendón-Anaya et al. (2017) for the split between a domesticated Andean genotype and a domesticated Mesoamerican genotype but differs greatly compared to the split between a wild genotype (G21245) from northern Peru/Ecuador and the aforementioned Andean and Mesoamerica genotypes (0.9 Mya). Based on our data, the divergence between the Mesoamerican and Andean genepools occurred much earlier ∼0.09 Mya, which is similar to earlier estimates of ∼0.11 Mya (Mamidi et al., 2013) and ∼0.087 Mya (Ariani et al., 2018). Based on all lines of evidence, we propose that two migratory events occurred, and both originated in Mesoamerica.

The first migratory event allowed the common bean to spread to northern Peru/Ecuador about 150,000 years ago. A more recent migration then gave rise the Andean genepool ∼90,000 years ago. Indeed, a single migration event would imply an initial adaptation to the equatorial environment and a subsequent adaptation to the negative latitude and high altitude. Furthermore, based on the same rationale, we suggest that the migration from Mesoamerica (Costa Rica) to northern Peru/Ecuador was facilitated by migratory birds flying over the Pacific Ocean. The second migration could involve either of two main bird migration routes from North to South America (Zimmer et al., 1938), one through the Gulf of Mexico and a longer one through Central America passing across the neck of Panama and then following the course of the Andes, or the west coast, or the northern coast eastward from Panama, or a diagonal course southeast through the Amazonian region. Migratory birds may therefore be responsible for several long-distance dispersal events (Remsen 1984; Sorte et al., 2016; Vianna et al., 2016) that led to the current distribution of the wild common bean. Differently, Ariani et al. (2018) also proposed at least three long-distance migratory events from the centre of origin in Mexico to southern Mesoamerica, northern Peru/Ecuador, and the southern Andes. However, our results indicate the occurrence of just two migration events.

Our attempt to reconstruct the phylogenetic history of the common bean using nuclear markers provides a clear example of the bias introduced by the use of markers from DNA regions subject to recombination. Indeed, one of the main assumptions during phylogenetic reconstruction is the absence of recombination. Ignoring crossovers, gene conversion, horizontal transfer and hybridization can lead to erroneous estimates that do not represent any of the probable evolutionary histories of the species. Previous studies in which phylogenesis was carried out using genome-wide markers placed PhI samples on the outermost branches (Rendón-Anaya et al., 2017, Ariani et al., 2018, Papa and Gepts, 2003). To overcome the consequences of recombination and especially crossover events, we restricted our analysis to SNPs located in centromeric regions, which are cold spots for recombination (Fernandes et al., 2019). However, the analysis of centromeric markers did not provide sufficient resolution to infer intraspecific phylogenetic relationships among *P. vulgaris* genotypes. Indeed, it was not possible to make, from different chromosomes, unique inferences about the derivation of PhI. Introgression, incomplete lineage sorting and gene conversion may have acted as non-reciprocal recombination events even in the absence of crossovers (Talbert and Henikoff, 2010), making the phylogenetic approach based on nuclear markers unsuitable for detecting close relationships, such as those between genepools of the same species. Nevertheless, the 11 trees (one for each centromeric region) revealed that even though the centromeres had slightly different topologies, the PhI sample visibly showed a relationship to the Mesoamerican genepool and specifically with accessions from Guatemala, Costa Rica and the valley of Oaxaca, the latter suggested as the presumed centre of domestication (Bitocchi et al., 2013; Rodriguez et al., 2016).

Our work is an example of phylogenetics applied to the evolutionary history of populations belonging to the same biological species, and in particular the three wild genepools of *P. vulgaris*. Our findings confirm the monophyletic and Mesoamerican origin of the common bean, but also provide a deeper understanding of the relationships between the three major wild genepools and their divergence events. In addition, we provide clear evidence of bias due to recombination events when using nuclear data to reconstruct phylogenetic trees. Our study therefore clarifies the intraspecific phylogeny of *P. vulgaris* and its origin in Mesoamerica.

## Materials and Methods

### Plant material, preparation of chloroplast/nuclear DNA libraries, and phaseolin extraction

Patterns of nucleotide variability were assessed across 97 *Phaseolus spp.* chloroplast DNA samples. These comprised 70 wild accessions of *P. vulgaris*, 22 of *P. coccineus*, three of *P. lunatus* and one of *P. acutifolius*, as well as one domesticated accession of *P. acutifolius*. The 70 *P. vulgaris* accessions covered the geographical distribution from northern Mexico to north-western Argentina and included all three genepools (44 Mesoamerican, 22 Andean and 4 northern Peru/Ecuador). Ten *P. vulgaris* accessions were selected from the original panel for resequencing and nuclear genome analysis. The selection was based on geographic criteria, guaranteeing the representation of Mesoamerica, Andes, and northern Peru/Ecuador, and on the haplogroups identified by analysing the chloroplast genome. Genomic DNA was extracted from the young leaves of single greenhouse-grown plants using the DNeasy Plant Mini Kit (Qiagen). Paired-end DNA libraries were constructed and sequenced from both ends using Illumina technology, with low coverage for the chloroplast genome samples and 10× coverage for the nuclear genome samples. Seeds were provided by the United States Department of Agriculture (USDA) Western Regional Plant Introduction Station and the International Center of Tropical Agriculture (CIAT) in Colombia. To verify the presence of the ancestral type I phaseolin protein, phaseolin was extracted from seed samples of representative accessions and enriched as described in the Supplementary Methods.

### Reference mapping and SNP calling in the chloroplast dataset

Quality control was applied to raw reads before and after trimming using FastQC (Andrews, 2010). Trimmomatic v0.38 (Bolger et al., 2014) was used to remove Illumina technical sequences and filter out low-quality reads. Reads ≥ 75 nucleotides in length with a minimum Q-value of 20 were retained. FastQscreen was used for contamination screening (Wingett et al., 2018) and chloroplast reads were retrieved by screening the *P. vulgaris* nuclear (G19833) and chloroplast (NC_009259) genomes using Bowtie2 with default settings (Langmead et al., 2012). The filtered reads were mapped to the chloroplast genome (NC_009259) using Bowtie2 with “local” settings, and the mapped reads were sorted and realigned with SAMtools v1.7 (Li et al., 2009). The read depth across the *P. vulgaris* chloroplast genome sequence was determined by using the BEDtools “genomecov” utility to find all uniquely mapping reads in the library (Quinlan et al., 2010).

SNPs in the *P. vulgaris* chloroplast genome sequence were called using the “mpileup” utility of BCFtools (Li et al., 2009). All VCF files were merged using BCFtools and uninformative SNPs (singletons) were filtered with VCFTOOLS (Danecek et al., 2011). The final set of SNPs was annotated with predicted functional effects using SnpEff (Cingolani et al., 2012). The VCF files were converted to Nexus and BAPS format using PGDSpider v2.1.1.5 (Lischer and Excoffier., 2012).

### Reference mapping and SNP calling in the nuclear dataset

SNPs in the *P. vulgaris* nuclear genome sequence were called using the sequence_handling pipeline (Hoffman et al., 2018) available at the Minnesota Supercomputing Institute, followed by quality control using FastQC. Adapters were removed using Scythe v.1.2.8 (https://github.com/vsbuffalo/scythe) with default parameters (e.g., prior contamination rate of 0.05). Sequences were aligned to the *P. vulgaris* reference genome (accession G19833, v.2.1) using BWA-MEM v0.7.17 with default parameters (Li, 2013). The resulting SAM files were sorted and de duplicated, and read groups were added with Picard v2.4.1 (http://broadinstitute.github.io/picard/). Haplotypes were called using GATK v4.1.2 (Poplin et al., 2017) with a nucleotide sequence diversity estimate (Watterson theta value) of θW = 0.001. The latter was estimated using ANGSD (Thorfinn et al., 2014) based on the samples and an ancestral sequence obtained by mapping *P. lunatus* reads to the *P. vulgaris* reference genome. The resulting gVCF files were used to jointly call SNPs on all samples. Hard-filtering was applied to increase the quality of the call-set (4Danecek et al., 2021). Indels, non-biallelic sites, low-quality sites (missingness ≥ 50%) and sites with minor allele frequencies ≤ 0.01 were filtered. Finally, singletons were removed from the final set of SNPs to avoid noisy signals due to long-branch attraction effects. SNPs were annotated with SnpEff as above.

### Genetic structure of the chloroplast dataset

The SNP dataset was clustered using BAPS v6.0 (Cheng et al., 2013). We chose a mixture model due to the high probability that all markers were linked. Pairwise identity-by-state distances were estimated among all 97 samples and among the *P. vulgaris* accessions using PLINK v1.90b52 (Purcell et al., 2007) and the results were graphically represented by MDS. Haplotype network analysis was carried out using PopART (Leigh and Bryant, 2015) with the TCS network (Clement et al., 2022).

### Assembly of chloroplast genomes

Full-length assembly was not possible for all 97 chloroplast genomes due to the low sequencing depth (Bedtools, “genomecov” utility). Therefore, we selected 31 *P. vulgaris* accessions based on coverage and geographical origin, representing all three gene pools: 19 Mesoamerican, 8 Andean, and 4 PhI. We also included four samples of *P. coccieneus*, one of *P. acutifolius* and one of *P. lunatus*. NOVOPlasty v3.2 (Dierckxsens et al., 2017) was used for de novo genome assembly, seeded with the *P. vulgaris* sequences matk, accd, psbh, rrn16 and rpl32 (GenBank accession no. NC_009259).

### Phylogenetic analysis

An ML tree based on the 37 chloroplast genome de novo assemblies was computed using RAxML v8.1.2 (Stamatakis, 2014) with the GTR substitution model, a bootstrap value of 10,000, and *P. lunatus* selected as the outgroup (Delgado-Salinas et al., 1999). The same analysis was applied to the concatenated SNPs, again with 10,000 bootstrap replicates. Trees were visualized using FigTree v1.4.4. An ML tree was also computed in RAxML for the nuclear dataset, only including those SNPs located in the pericentromeric regions. The SNPs were concatenated, and an individual ML tree was constructed for each of the 11 chromosomes, with 10,000 bootstrap replicates.

### Dating the divergence events

Molecular clock analysis was applied to the 37 chloroplast genome *de novo* assemblies. These were aligned using MAFFT (Katoh et al., 2019) to three *Vigna spp.* chloroplast genomes in GenBank, namely *V. radiata* (NC_013843), *V. unguiculata* (NC_018051) and *V. angularis* (NC_021091), followed by Bayesian analysis in BEAST v2.6.2 (Bouckaert et al., 2014). The *BEAST method was used to produce the XML file and the coalescent simulation was initiated by applying a relaxed log-normal molecular clock with a general time reversible model. The tree was calibrated using the divergence reported between *P. coccineus* and *Vigna spp.* (Lavin et al., 2005), which is μ = 1.23×10-3 substitutions per site per year. The Monte Carlo Markov chain was set to 100,000,000 and two independent runs were combined.

## Acknowledgments

This work was supported by grants from the Italian Government (MIUR; Grant number 20177RL4KL, PRIN – PARDOM Project 2017), the Università Politecnica delle Marche (2020-2022) and INCREASE project, funded from the European Union Horizon 2020 Research and Innovation program under grant agreement no. 862862.

## Author Contributions

G. F., E. Bitocchi, and R.P. designed research; G. F., T. R. G., N. D. A., E. Bitocchi, and R.P. analyzed data; G. F. and R.P. wrote the paper; D. C., S. C., G. C., F. S., E. Bellucci, V. D. V., L. N., A. P., M. R., L. V., A. B., M. D., contributed to the research activity and the editing of the article; D. C., S. C., M. R., L. V. and M.D. conducted the sequencing. We declare no conflict of interest.

## Data availability

The raw sequence reads generated and analysed during this study are available in the Sequence Read Archive (SRA) of the National Center for Biotechnology Information (NCBI) with the BioProject number PRJNA910538.

The bioinformatics pipelines used in this study are available in https://github.com/giuliafrascarelli/Common_bean.

## References

Andrews S. FastQC: A Quality Control Tool for High Throughput Sequence Data [Online] (2010). Available online at: http://www.bioinformatics.babraham.ac.uk/projects/fastqc/

Ariani A., J. C. B. Mier Teran, P. Gepts. “Spatial and Temporal Scales of Range Expansion in Wild Phaseolus Vulgaris.” Molecular Biology and Evolution 35(1):119–31 (2018). doi: 10.1093/molbev/msx273.

Avise J. C. Phylogeography: retrospect and prospect. Journal of Biogeography. Vol. 36, 3–15 (2009). 10.1111/j.1365-2699.2008.02032.x

Avise J.C., J. Arnold, R.M. Ball, Jr E. Bermingham, T. Lamb, J.E. Neigel, C.A. Reeb & N.C. Saunders (1987) Intra-specific phylogeography: the mitochondrial DNA bridgebetween population genetics and systematics. Annual Reviewof Ecology and Systematics,Vol. 18,489–522.

Bitocchi E., E. Bellucci, A. Giardini, D. Rau, M. Rodriguez, E. Biagetti, A. Carboni, P. Gepts, L. Nanni, R. Papa, and G. Attene. “Molecular Analysis of the Parallel Domestication of the Common Bean (Phaseolus Vulgaris) in Mesoamerica and the Andes.” New Phytologist 197(1):300–313 (2013). doi: 10.1111/j.1469-8137.2012.04377.x.

Bitocchi E., L. Nanni, E. Bellucci, M. Rossi, A. Giardini,, D. Rau, M. Rodriguez, E. Biagetti, R. Santilocchi, P. Spagnoletti Zeuli, T. Gioia, G. Logozzo, G. Attene, L. Nanni, and R. Papa. “Mesoamerican Origin of the Common Bean (Phaseolus Vulgaris L.) Is Revealed by Sequence Data.” Proceedings of the National Academy of Sciences of the United States of America 109(14) (2012). doi: 10.1073/pnas.1108973109.

Bolger A. M., M. Lohse, and B. Usadel. “Trimmomatic: A Flexible Trimmer for Illumina Sequence Data.” Bioinformatics 30(15):2114–20 (2014). doi: 10.1093/bioinformatics/btu170.

Bouckaert R., J. Heled, D. Kühnert, T. Vaughan, C. Hsi Wu, D. Xie, M. A. Suchard, A. Rambaut, and A. J. Drummond. “BEAST 2: A Software Platform for Bayesian Evolutionary Analysis.” PLoS Computational Biology 10(4) (2014). doi: 10.1371/journal.pcbi.1003537.

Cavalli-Sforza L. L., and A. W. F. Edwards. Phylogenetic Analysis: Models and Estimation Procedures (1966). doi: 10.1111/j.1558-5646.1967.tb03411.x.

Cheng L., T. R. Connor, J. Sirén, D. M. Aanensen, J. Corander. Hierarchical and Spatially Explicit Clustering of DNA Sequences with BAPS Software, Molecular Biology and Evolution, Vol. 30. Issue 5. Pages 1224–1228 (2013), 10.1093/molbev/mst028

Cingolani P., A. Platts, L. L. Wang, M. Coon, T. Nguyen et al. “A Program for Annotating and Predicting the Effects of Single Nucleotide Polymorphisms, SnpEff: SNPs in the Genome of Drosophila Melanogaster Strain W1118; Iso-2; Iso-3.” Fly 6(2):80–92 (2012). doi: 10.4161/fly.19695.

Clement M., Q. Snell, P. Walker, D. Posada, and K. Crandall. TCS: Estimating Gene Genealogies. In Parallel and Distributed Processing Symposium, International (Vol. 3, pp. 0184–0184). IEEE Computer Society (2002).

Danecek P., A. Auton, G. Abecasis, C. A. Albers, E. Banks, M. A. DePristo, R. E. Handsaker, G. Lunter, G. T. Marth, S. T. Sherry, G. McVean, and R. Durbin“The Variant Call Format and VCFtools.” Bioinformatics 27(15):2156–58 (2011). doi: 10.1093/bioinformatics/btr330.

Danecek P., J. K. Bonfield, J. Liddle, J. Marshall, V. Ohan, M. O. Pollard, A. Whitwham, T. Keane, S. A. McCarthy, R. M. Davies, and H. Li “Twelve Years of SAMtools and BCFtools.” GigaScience 10(2) (2021). doi: 10.1093/gigascience/giab008.

Debouck D.G., 1986. Phaseolus germplasm collection in Cajamarca and Amazonas, Peru. International Board for Plant Genetic Resources, Rome, Italy. AGPG/IBPGR: 86/161. Mimeographed, 36 p.

Delgado-Salinas A., T. Turley, A. Richman, and M. Lavin. “Phylogenetic Analysis of the Cultivated and Wild Species of Phaseolus (Fabaceae).” Systematic Botany 24(3):438–60 (1999). doi: 10.2307/2419699.

Desiderio F., E. Bitocchi, E. Bellucci, D. Rau, M. Rodriguez, G. Attene, R. Papa, and L. Nanni “Chloroplast Microsatellite Diversity in Phaseolus Vulgaris.” Frontiers in Plant Science 3(JAN) (2013). doi: 10.3389/fpls.2012.00312.

Dierckxsens N., P. Mardulyn, and G. Smits. “NOVOPlasty: De Novo Assembly of Organelle Genomes from Whole Genome Data.” Nucleic Acids Research 45(4) (2017). doi: 10.1093/nar/gkw955.

Fernandes J. B., P. Wlodzimierz, I. R. Henderson. Meiotic recombination within plant centromeres. Current Opinion in Plant Biology. Vol. 48, 26–35. ISSN 1369-5266 (2019), 10.1016/j.pbi.2019.02.008.

Gepts P. Biochemical Evidence Bearing on the Domestication of Phaseolus (Fabaceae) Beans I. Vol. 44 (1990).

Hoffman P. J., S. R. Wyant, T. J. Y. Kono, P. L. Morrell. sequence_handling: A pipeline to automate sequence workflows. Zenodo (2018). 10.5281/zenodo.1257692.

Kami J., V. Becerra Velasquez, D. G. Debouck, P. Gepts, and R. W. Allard. Identification of Presumed Ancestral DNA Sequences of Phaseolin in Phaseolus Vulgaris (Molecular Evolution/Seed Protein/Crop Evolution/Tandem Repeat/Polymerase Chain Reaction) Communicated By. Vol. 92. (1995) doi: 10.1073/pnas.92.4.1101.

Katoh K., J. Rozewicki, K. D. Yamada. MAFFT online service: multiple sequence alignment, interactive sequence choice and visualization. Brief Bioinform. (2019) Jul 19;20(4):1160–1166. doi: 10.1093/bib/bbx108. PMID: 28968734; PMCID: PMC6781576.

Kwak M., J. A. Kami, and P. Gepts. “The Putative Mesoamerican Domestication Center of Phaseolus Vulgaris Is Located in the Lerma-Santiago Basin of Mexico.” Crop Science 49(2):554–63 (2009). doi: 10.2135/cropsci2008.07.0421.

Langmead B., and S. L. Salzberg. “Fast Gapped-Read Alignment with Bowtie 2.” Nature Methods 9(4):357–59 (2012). doi: 10.1038/nmeth.1923.

Lavin M., P. S. Herendeen, and M. F. Wojciechowski. “Evolutionary Rates Analysis of Leguminosae Implicates a Rapid Diversification of Lineages during the Tertiary.” Systematic Biology 54(4):575–94 (2005). doi: 10.1080/10635150590947131.

Leigh J. W., and D. Bryant. “POPART: Full-Feature Software for Haplotype Network Construction.” Methods in Ecology and Evolution 6(9):1110–16 (2015). doi: 10.1111/2041-210X.12410.

Li H., Aligning sequence reads, clone sequences and assembly contigs with BWA-MEM. arXiv preprint arXiv:1303.3997 (2013)

Li H., B. Handsaker, A. Wysoker, T. Fennell, J. Ruan, N. Homer, G. Marth, G. Abecasis, and R. Durbin. “The Sequence Alignment/Map Format and SAMtools.” Bioinformatics 25(16):2078–79 (2009). doi: 10.1093/bioinformatics/btp352.

Lischer H. E. L. and Excoffier L. PGDSpider: An automated data conversion tool for connecting population genetics and genomics programs. Bioinformatics, 28, 298– 299 (2012). 10.1093/bioinformatics/btr642.

Mamidi S., M. Rossi, S. M. Moghaddam, D. Annam, R. Lee, R. Papa, and P. E. McClean. “Demographic Factors Shaped Diversity in the Two Gene Pools of Wild Common Bean Phaseolus Vulgaris L.” Heredity 110(3):267–76 (2013). doi: 10.1038/hdy.2012.82.

Meyer R.S., A.E. DuVal and H.R. Jensen (2012), Patterns and processes in crop domestication: an historical review and quantitative analysis of 203 global food crops. New Phytologist, 196: 29–48 (2012). 10.1111/j.1469-8137.2012.04253.x.

Nordborg M., S. Tavaré. Linkage disequilibrium: what history has to tell us. TRENDS in Genetics Vol.18 No.2. (2002) 10.1016/S0168-9525(02)02557-X

Papa R., P. Gepts. Asymmetry of gene flow and differential geographical structure of molecular diversity in wild and domesticated common bean (Phaseolus vulgaris L.) from Mesoamerica. Thero Appl Genet 106, 239–250 (2003). 10.1007/s00122-002-1085-z.

Poplin R., V. Ruano-Rubio, M. A. DePristo, T. J. Fennell, M. O. Carneiro, G. A. van der Auwera, D. E. Kling, L. D. Gauthier, A. Levy-Moonshine, D. Roazen, K. Shakir, J. Thibault, S. Chandran, C. Whelan, M. Lek, S. Gabriel, M. J. Daly, B. Neale, D. G. MacArthur, and E. Banks.“Scaling Accurate Genetic Variant Discovery to Tens of Thousands of Samples.” BioRxiv (2017). doi: 10.1101/201178.

Purcell S., B. Neale, K. Todd-Brown, L. Thomas, M.A.R. Ferreira, D. Bender, J. Maller, P. Sklar, P. I. W. de Bakker, M. J. Daly, P. C. Sham. PLINK: a tool set for whole-genome association and population-based linkage analyses. The American Journal of Human Genetics, 81, 559– 575 (2007).

Quinlan A. R., and I. M. Hall. “BEDTools: A Flexible Suite of Utilities for Comparing Genomic Features.” Bioinformatics 26 (6): 841–42 (2010). doi: 10.1093/bioinformatics/btq033.

Remsen V. High incidence of “leapfrog” pattern of geographicvariation in Andean birds: implications for the speciation process. Science 224 (4645):171–173 (1984).

Rendón-Anaya M., A. Herrera-Estrella, P. Gepts, A. Delgado-Salinas. A new species of Phaseolus (Leguminosae, Papilionoideae) sister to Phaseolus vulgaris, the common bean. Phytotaxa, 313(3), 259–266 (2017).

Rendón-Anaya M., J.M. Montero-Vargas, S. Saburido-Álvarez, A. Vlasova, S. Capella-Gutierrez, J. Juan Ordaz-Ortiz, O. M. Aguilar, R. P. Vianello-Brondani, M. Santalla, L. Delaye, T. Gabaldón, P. Gepts, R. Winkler, R. Guigó, A. Delgado-Salinas, and A. Herrera-Estrella. 2017. “Genomic History of the Origin and Domestication of Common Bean Unveils Its Closest Sister Species.” Genome Biology 18(1). doi: 10.1186/s13059-017-1190-6.

Rodriguez M., D. Rau, E. Bitocchi, E. Bellucci, E. Biagetti, A. Carboni, P. Gepts, L. Nanni, R. Papa, and G. Attene. “Landscape Genetics, Adaptive Diversity and Population Structure in Phaseolus Vulgaris.” New Phytologist 209(4):1781–94 (2016). doi: 10.1111/nph.13713.

Rossi M., E. Bitocchi, E. Bellucci, L. Nanni, D. Rau,, G. Attene, and R. Papa. “Linkage Disequilibrium and Population Structure in Wild and Domesticated Populations of Phaseolus Vulgaris L.” Evolutionary Applications 2(4):504–22 (2009). doi: 10.1111/j.1752-4571.2009.00082.x.

Schierup M. H., and J. Hein. Consequences of Recombination on Traditional Phylogenetic Analysis. Genetics, Vol. 156 (2, 1) (2000): 879–891/. doi: 10.1093/genetics/156.2.879

Sorte F., D. Fink, W. M. Hochachka, S. Kelling. Convergence ofbroad-scale migration strategies in terrestrial birds. Proc R Soc B283:20152588 (2016).

Stamatakis A. “RAxML Version 8: A Tool for Phylogenetic Analysis and Post-Analysis of Large Phylogenies.” Bioinformatics 30(9):1312–13 (2014). doi: 10.1093/bioinformatics/btu033.

Talbert P. B., S. Henikoff. Centromeres Convert but Don’t Cross. PLOS Biology 8(3): e1000326 (2010). 10.1371/journal.pbio.1000326.

Thorfinn S. K., A. Albrechtsen, and R. Nielsen. ANGSD: Analysis of Next Generation Sequencing Data. BMC Bioinformatics 15, 356 (2014). Doi: 10.1186/s12859-014-0356-4.

Toro C., O. J. M. Tohme, and D. G. Debouck. Wild Bean (Phaseolus Vulgaris L.): Description and Distribution. Vol. 181. CIAT (1990)

Vianna D.S, L. Gangoso, W. Bouten, J. Figuerola. Overseas seed dis-persal by migratory birds.Proc Biol Sci. 283(1822):20152406 (2016).

Wingett S. W., and S. Andrews. “Fastq Screen: A Tool for Multi-Genome Mapping and Quality Control.” F1000Research 7 (2018). doi: 10.12688/f1000research.15931.1.

Zimmer J. T. Notes on migrations of South American birds. Auk55: 405–410 (1938).

