## Supplementary File for "The evolutionary history of the common bean (*Phaseolus vulgaris*) revealed by chloroplast and nuclear genomes"

### Supplementary Materials

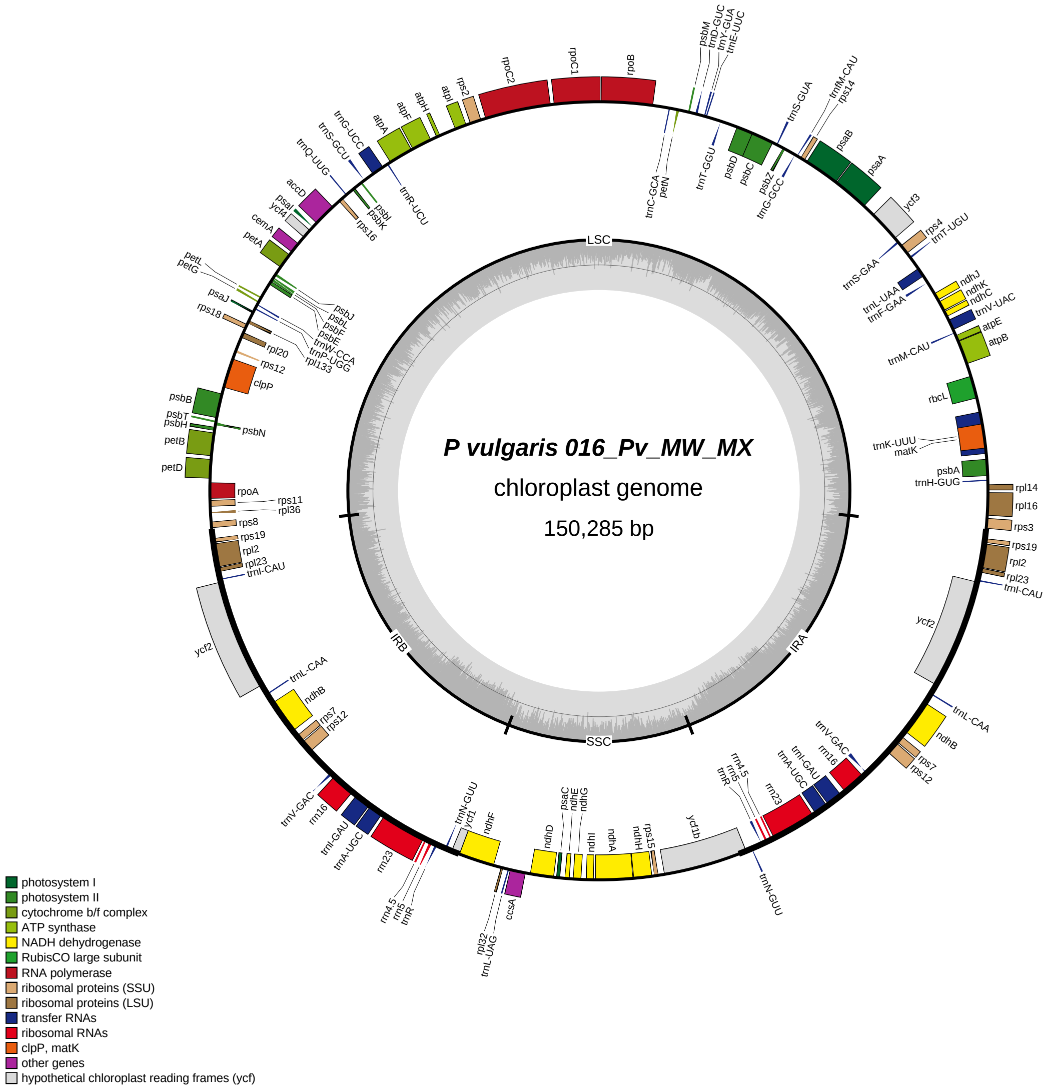


Supplementary_Figure_1: Map of the chloroplast genome of the accessions 016_Pv_MW_MX (*Phaseolus vulgaris*). Genes inside of the outer circle are transcribed in the clockwise direction, while those outsides are transcribed in the counterclockwise direction. Different color codes represent genes belonging to various functional groups. The circle inside represents GC content graph with the 50% threshold. The inverted repeat, large single-copy, and small single-copy regions are denoted by IR, LSC, and SSC, respectively.

**Comparative analysis**

To visualize the differences among the assembled chloroplast genomes, a multiple alignment was built in mVISTA (Stanford University, Stanford CA, USA) in LAGAN (Limited Area Global Alignment of Nucleotides) mode using *P. vulgaris* published plastome and its annotation as reference (NC_009259). An additional alignment was carried out using the MAUVE alignment software (Darling et al. 2004). This alignment was performed on a subset of plastomes, chosen based on the gene content differences revealed by the annotation, using the progressive MAUVE option.

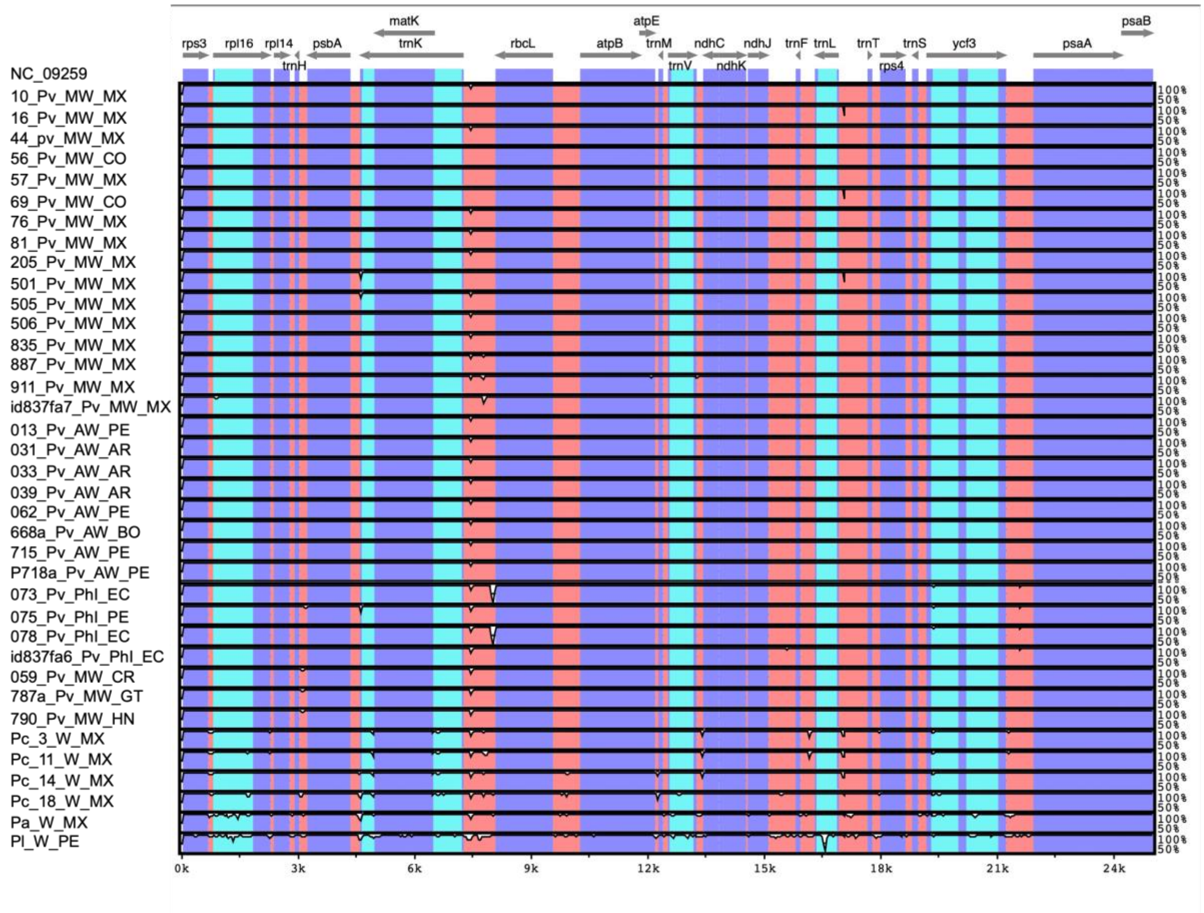

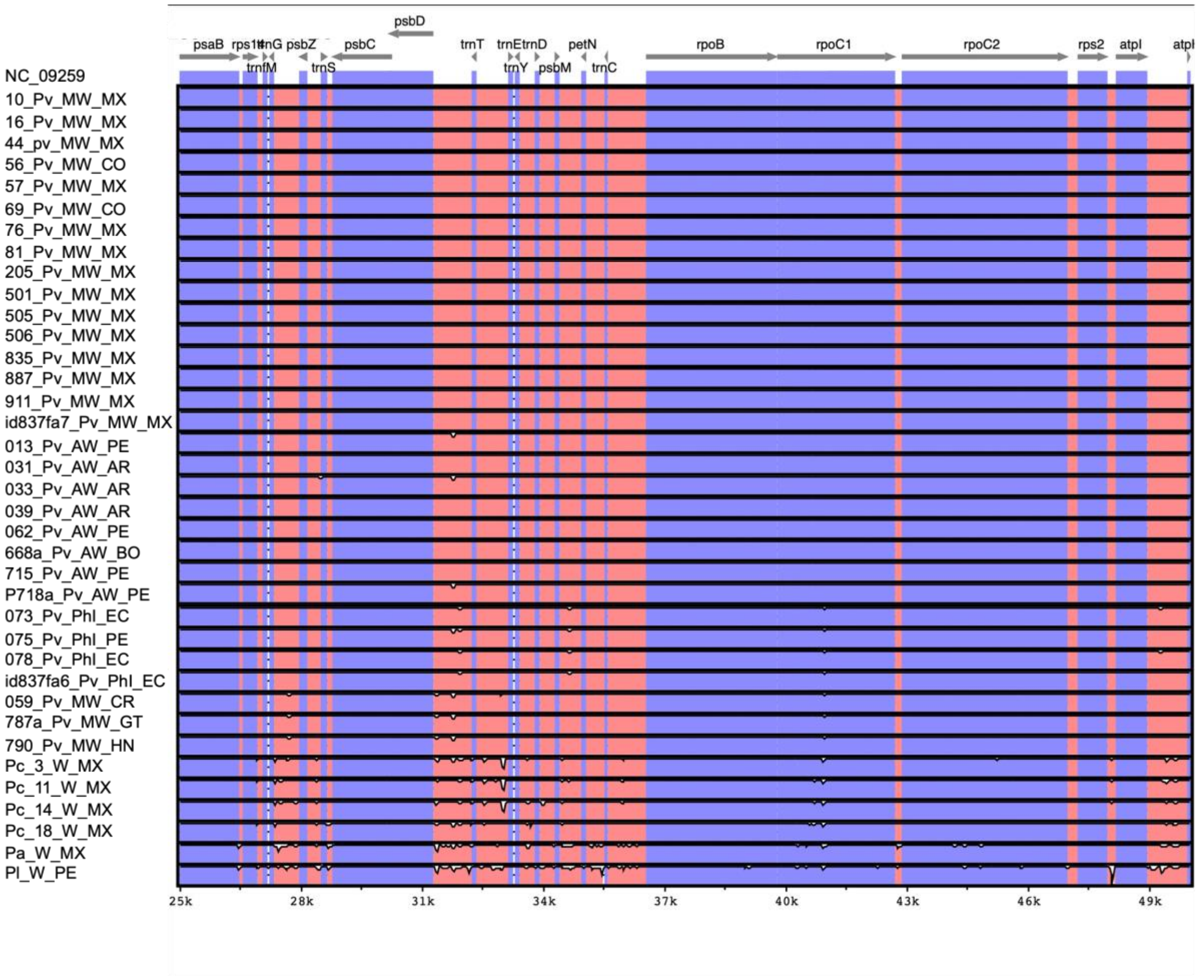
psbD
psaB
rps14
trnG
psbZ
psbC
trnfM
trnS
trnT
trnE
trnD
trnY
psbM
petN
trnC
rpoB
rpoC1
rpoC2
rps2
atpI
atpI
NC_09259
10_Pv_MW_MX
16_Pv_MW_MX
44_pv_MW_MX
56_Pv_MW_CO
57_Pv_MW_MX
69_Pv_MW_CO
76_Pv_MW_MX
81_Pv_MW_MX
205_Pv_MW_MX
501_Pv_MW_MX
505_Pv_MW_MX
506_Pv_MW_MX
835_Pv_MW_MX
887_Pv_MW_MX
911_Pv_MW_MX
id837fa7_Pv_MW_MX
013_Pv_AW_PE
031_Pv_AW_AR
033_Pv_AW_AR
039_Pv_AW_AR
062_Pv_AW_PE
668a_Pv_AW_BO
715_Pv_AW_PE
P718a_Pv_AW_PE
073_Pv_Phl_EC
075_Pv_Phl_PE
078_Pv_Phl_EC
id837fa6_Pv_Phl_EC
059_Pv_MW_CR
787a_Pv_MW_GT
790_Pv_MW_HN
Pc_3_W_MX
Pc_11_W_MX
Pc_14_W_MX
Pc_18_W_MX
Pa_W_MX
Pl_W_PE
25k
28k
31k
34k
37k
40k
43k
46k
49k

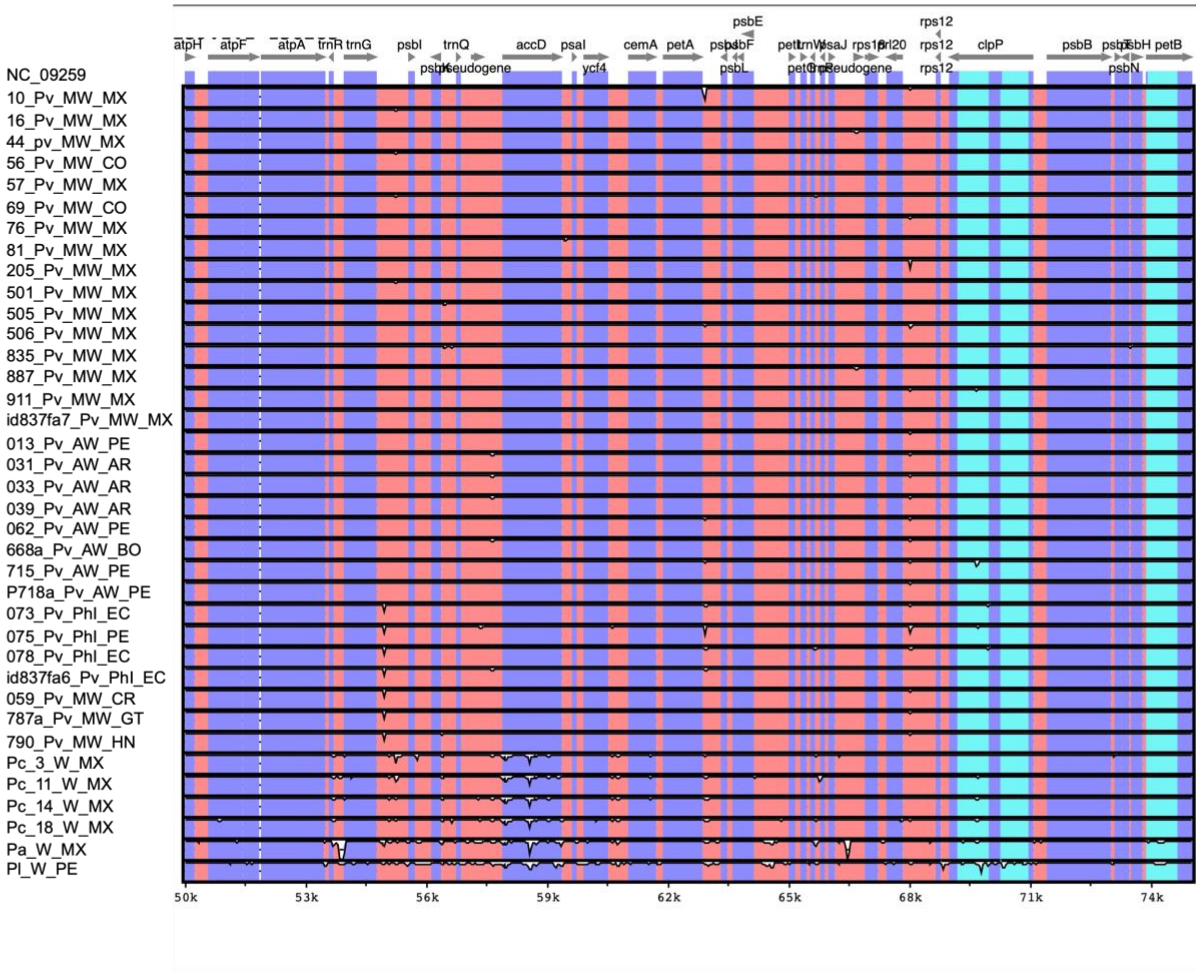
atpH
atpF
atpA
trnR
trnG
psbI
psbK
trnQ
pseudogene
accD
psaI
ycf4
cemA
petA
psbE
psbF
psbL
petL
petG
psaJ
rps18
rpl20
rps12
clpP
psbB
psbN
psbH
petB
NC_09259
10_Pv_MW_MX
16_Pv_MW_MX
44_pv_MW_MX
56_Pv_MW_CO
57_Pv_MW_MX
69_Pv_MW_CO
76_Pv_MW_MX
81_Pv_MW_MX
205_Pv_MW_MX
501_Pv_MW_MX
505_Pv_MW_MX
506_Pv_MW_MX
835_Pv_MW_MX
887_Pv_MW_MX
911_Pv_MW_MX
id837fa7_Pv_MW_MX
013_Pv_AW_PE
031_Pv_AW_AR
033_Pv_AW_AR
039_Pv_AW_AR
062_Pv_AW_PE
668a_Pv_AW_BO
715_Pv_AW_PE
P718a_Pv_AW_PE
073_Pv_PhI_EC
075_Pv_PhI_PE
078_Pv_PhI_EC
id837fa6_Pv_PhI_EC
059_Pv_MW_CR
787a_Pv_MW_GT
790_Pv_MW_HN
Pc_3_W_MX
Pc_11_W_MX
Pc_14_W_MX
Pc_18_W_MX
Pa_W_MX
Pl_W_PE
50k
53k
56k
59k
62k
65k
68k
71k
74k

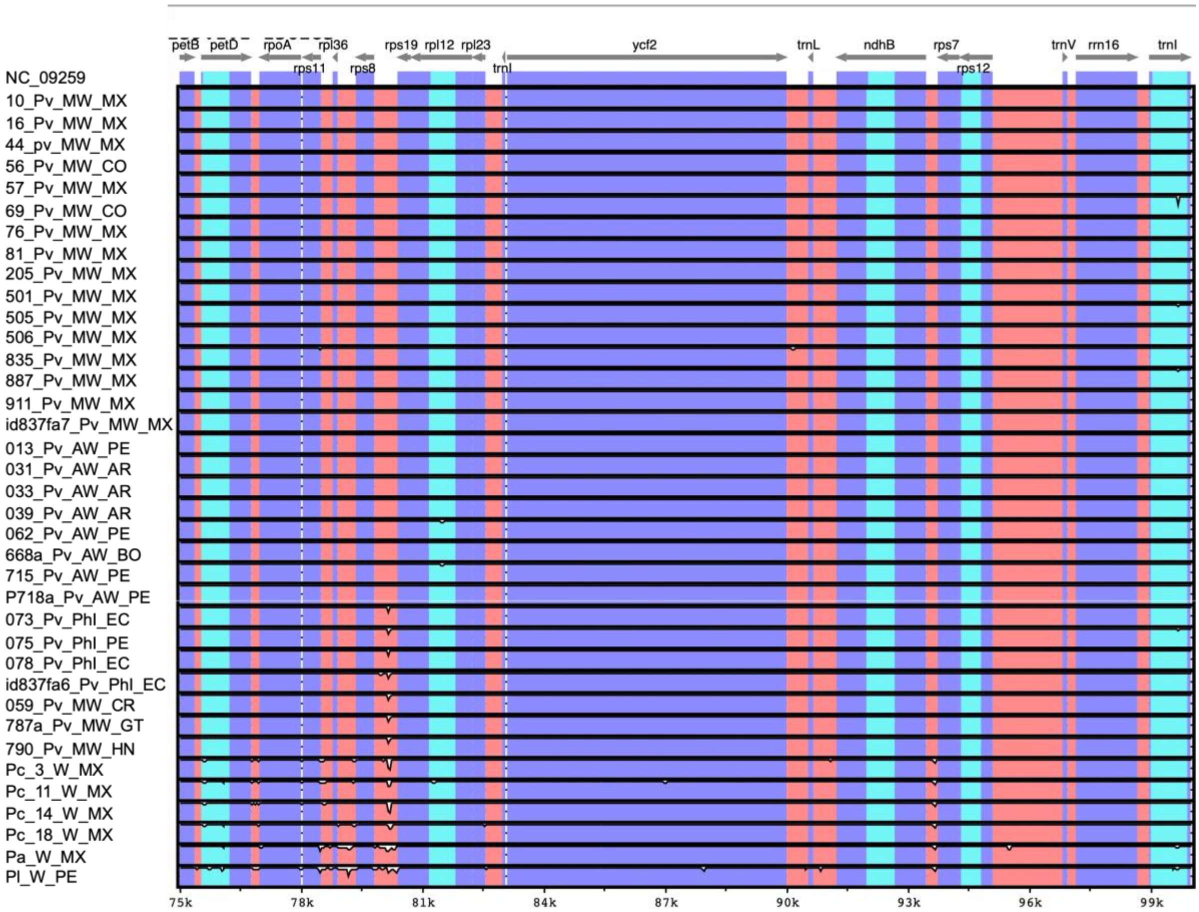

petB
petD
rpoA
rpl36
rps11
rps8
rps19
rpl12
rpl23
trnI
ycf2
trnL
ndhB
rps7
rps12
trnV
rrn16
trnI
NC_09259
10_Pv_MW_MX
16_Pv_MW_MX
44_pv_MW_MX
56_Pv_MW_CO
57_Pv_MW_MX
69_Pv_MW_CO
76_Pv_MW_MX
81_Pv_MW_MX
205_Pv_MW_MX
501_Pv_MW_MX
505_Pv_MW_MX
506_Pv_MW_MX
835_Pv_MW_MX
887_Pv_MW_MX
911_Pv_MW_MX
id837fa7_Pv_MW_MX
013_Pv_AW_PE
031_Pv_AW_AR
033_Pv_AW_AR
039_Pv_AW_AR
062_Pv_AW_PE
668a_Pv_AW_BO
715_Pv_AW_PE
P718a_Pv_AW_PE
073_Pv_Phl_EC
075_Pv_Phl_PE
078_Pv_Phl_EC
id837fa6_Pv_Phl_EC
059_Pv_MW_CR
787a_Pv_MW_GT
790_Pv_MW_HN
Pc_3_W_MX
Pc_11_W_MX
Pc_14_W_MX
Pc_18_W_MX
Pa_W_MX
Pl_W_PE
75k
78k
81k
84k
87k
90k
93k
96k
99k

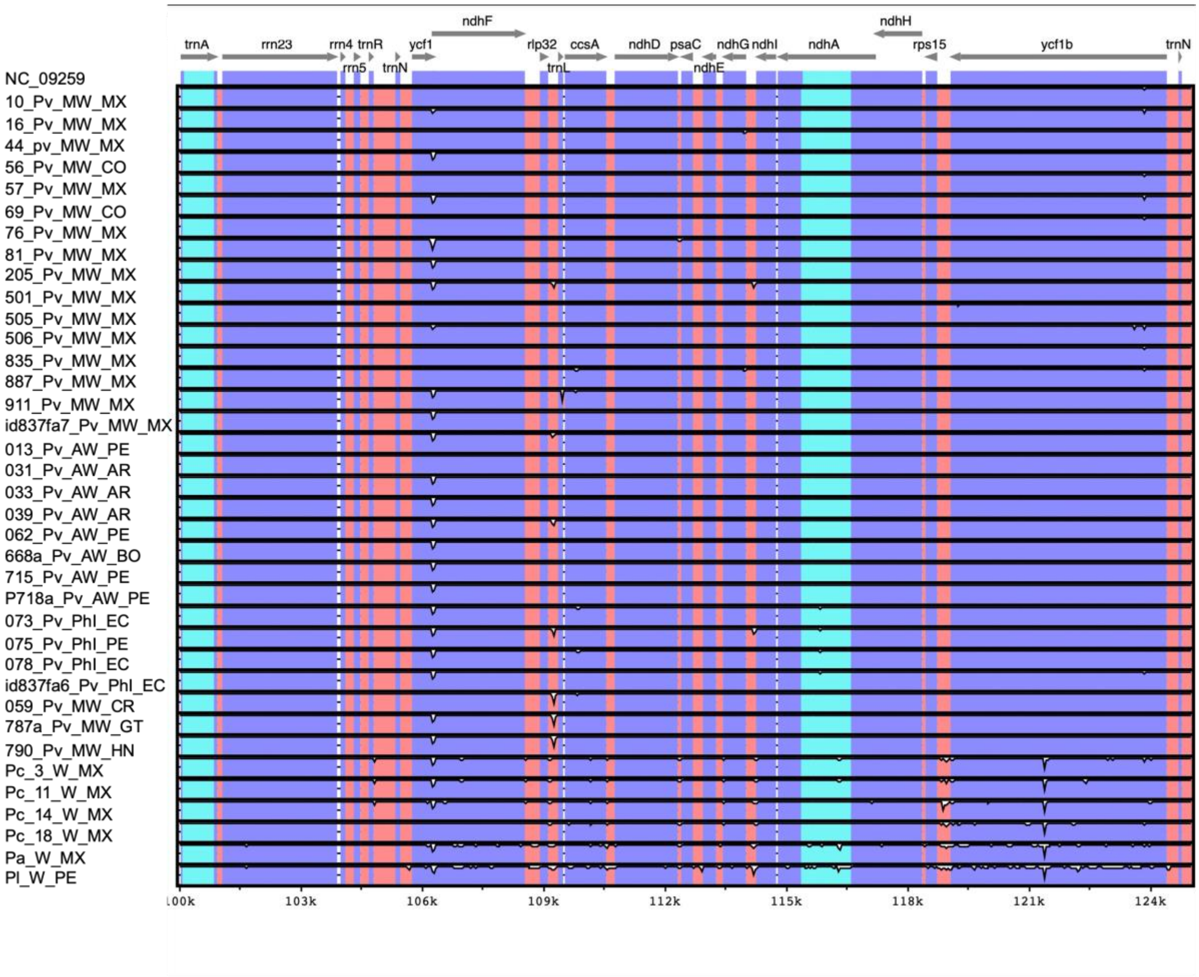
ndhF
ndhH
trnA
rrn23
rrn4
trnR
ycf1
rlp32
ccsA
ndhD
psaC
ndhG
ndhI
ndhA
rps15
ycf1b
trnN
rrn5
trnN
trnL
ndhE
NC_09259
10_Pv_MW_MX
16_Pv_MW_MX
44_pv_MW_MX
56_Pv_MW_CO
57_Pv_MW_MX
69_Pv_MW_CO
76_Pv_MW_MX
81_Pv_MW_MX
205_Pv_MW_MX
501_Pv_MW_MX
505_Pv_MW_MX
506_Pv_MW_MX
835_Pv_MW_MX
887_Pv_MW_MX
911_Pv_MW_MX
id837fa7_Pv_MW_MX
013_Pv_AW_PE
031_Pv_AW_AR
033_Pv_AW_AR
039_Pv_AW_AR
062_Pv_AW_PE
668a_Pv_AW_BO
715_Pv_AW_PE
P718a_Pv_AW_PE
073_Pv_Phl_EC
075_Pv_Phl_PE
078_Pv_Phl_EC
id837fa6_Pv_Phl_EC
059_Pv_MW_CR
787a_Pv_MW_GT
790_Pv_MW_HN
Pc_3_W_MX
Pc_11_W_MX
Pc_14_W_MX
Pc_18_W_MX
Pa_W_MX
Pl_W_PE
100k
103k
106k
109k
112k
115k
118k
121k
124k

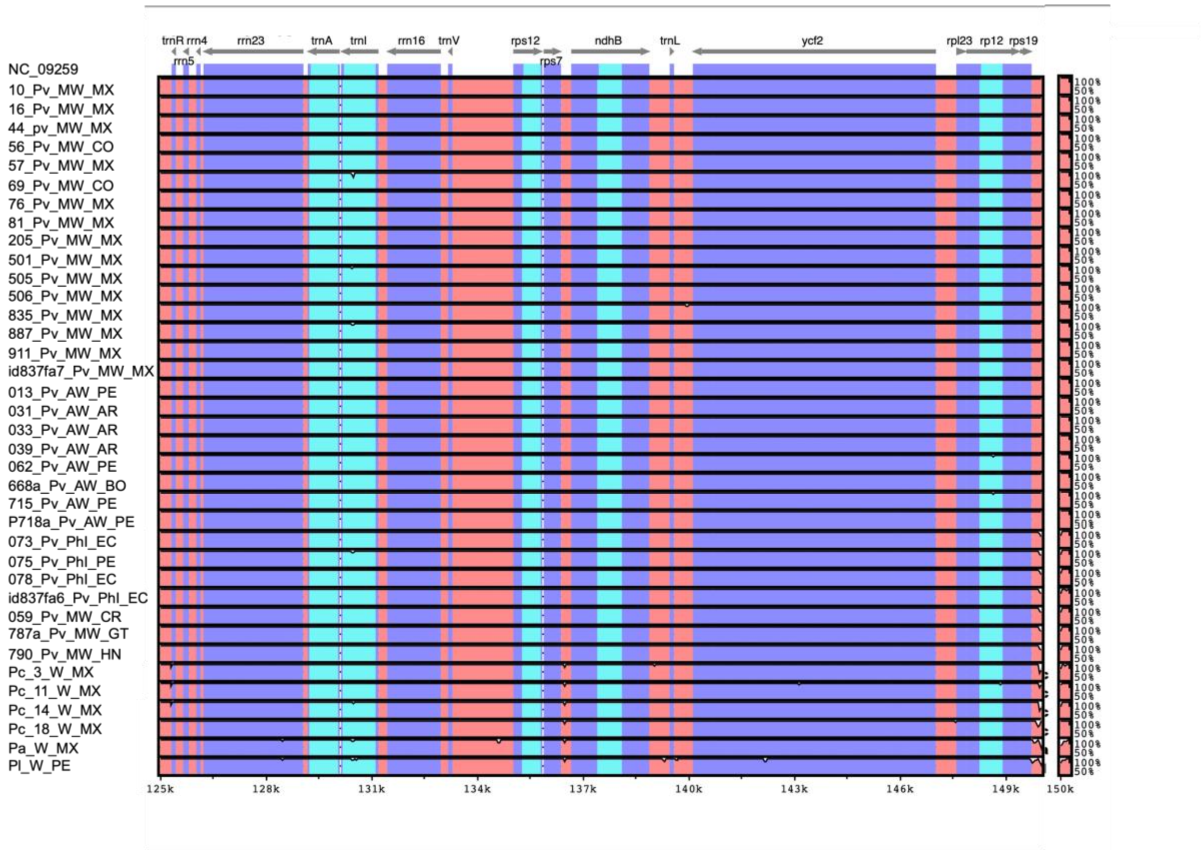


Supplementary_Figure_2: Sequence similarity plot by using mVISTA, among 37 de-novo assembled chloroplast genomes and NC_09259 as reference. In the y-axis percentage of sequence identity was shown between 50% and 100%. Transcriptional orientations of genes were assigned by grey arrows. Red bars represented non-coding sequences (NCS), purples bars represented exons and light blue bars represented untranslated regions (UTRs). Genomic differences were shown as white peaks.

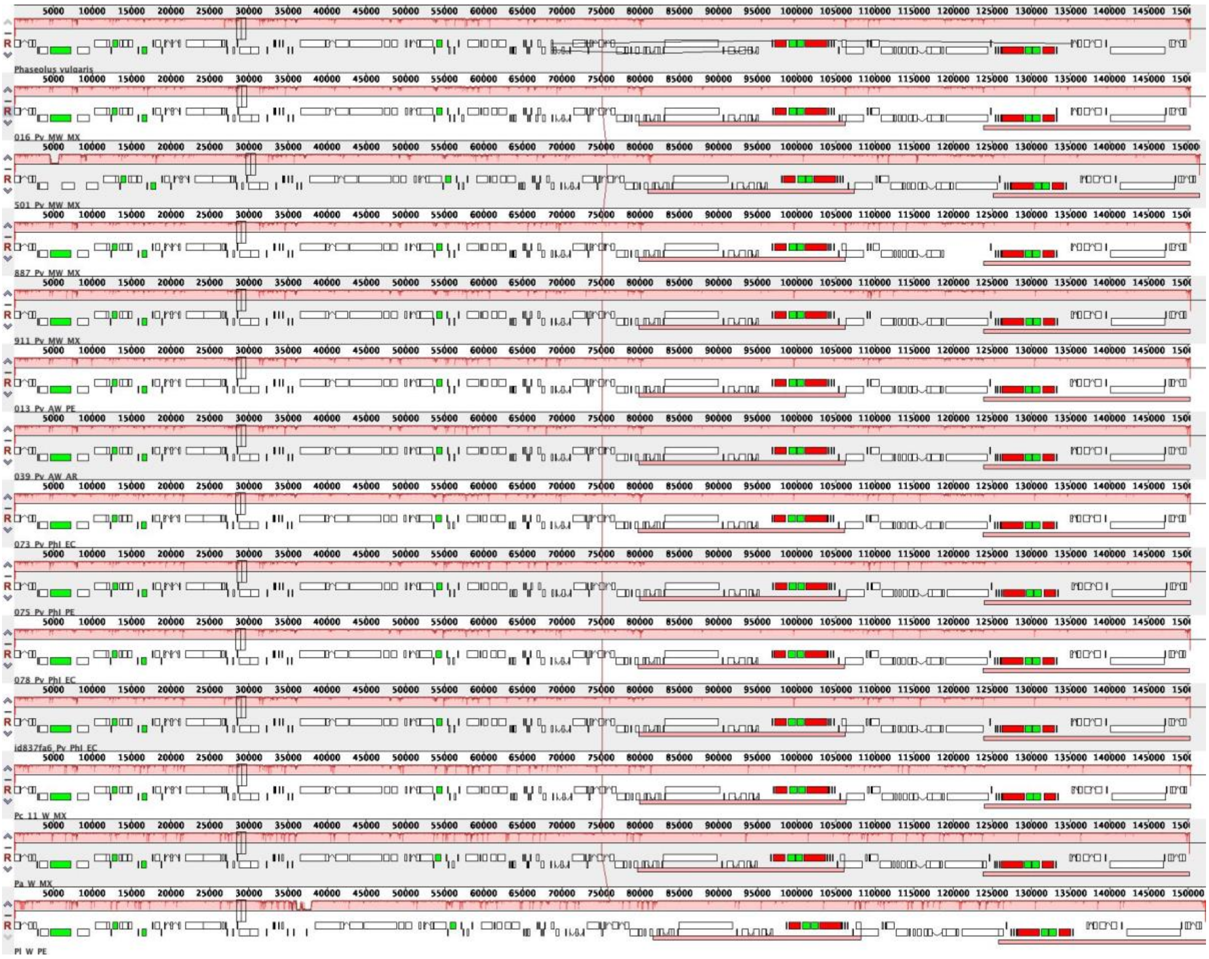

Supplementary_Figure_3: MAUVE alignment showing the gene order and homology between the reference chloroplast genome of *Phaseolus vulgaris* (NC_09259) fifteen *P. vulgaris* accessions (16_Pv_MW_MX, 501_Pv_MW_MX, 887_Pv_MW_MX, 911_Pv_M_MX, 013_Pv_AW_PE0, 39_Pv_AW_AR,54_Pv_PhI_PE, 073_Pv_PhI_EC, 075_Pv_PhI_PE, 078_Pv_PhI_EC), *P. coccineus* (Pc_11_W_MX), *P. acutifolius* (Pa_W_MX), *P. lunatus* (Pl_W_PE). Locally Collinear Blocks (LCBs) include the histograms that show sequence identity with peaks. Protein coding genes, rRNA genes, tRNA genes and intron containing tRNA genes are marked with block in white, red, black and green colors, respectively.

The genome variation among all *de-novo* assembled chloroplast genomes was analyzed with the online tool mVISTA unsing *P. vulgaris* (NC_09259) as reference. The mVISTA identity plot did not reveal meaningful differences between the *P. vulgaris* plastomes, with the exception of a small deletion in the intergenic region between *trnK* and *rbcL* gene, which was found in two PhI samples (i.e., 073_Pv_PhI_EC and 078_Pv_PhI_EC). In addition, a deletion (over 300 pb) was identified in the plastome of the *P. acutifolius* accession (i.e., Pa_W_MX) in the intergenic region between the *trnR* and *trnG* genes (Supprementary_Figure_2).

To identify the gene order and organization, a subset of 15 plastomes was aligned with MAUVE (Supplementary_Figure_3). Most of the regions were conserved among all plastomes and no rearrangements of gene order was detected.

**Analysis of nucleotide diversity**

In order to determine nucleotide diversity in the 37 *de-novo* assembled plastomes a multi-sequence alignment (MSA) was performed using MAFFT (Katoh et al., 2019) with default parameters. The MSA was used as input for DNAsp (Rozas et al., 2017) and nucleotide diversity was calculated. A sliding window of 200 bp with 50 bp step size was used to summarize the diversity statistics for visualization.

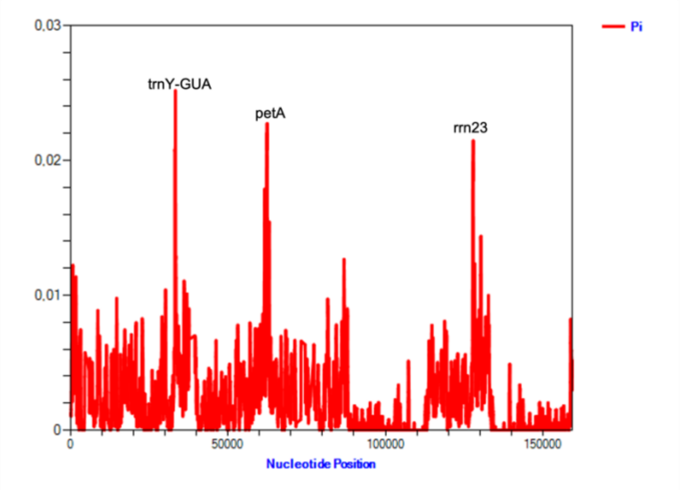


Supplementary_Figure_4: Sliding window analysis among the whole chloroplast genome of 37 de-novo assembled plastomes, including *P. vulgaris, P. coccineus, P. acutifolius, P. lunatus.* Regions with higher nucleotide variability are indicated.

The nucleotide diversity was investigated with DNAsp software. The multi sequence alignment revealed 145133 monomorphic sites 3586 polymorphic sites of which 1254 were defined as informative. A sliding window analysis was performed to calculate the nucleotide variability (Pi) across the 37 *de-novo* chloroplast genomes. Even though, the results showed a high sequence similarity, three divergent hot spots were detected (Pi>0.02): *trnY-GUA*, *petA*, *rrn23* (Supplementary_Figure_4).

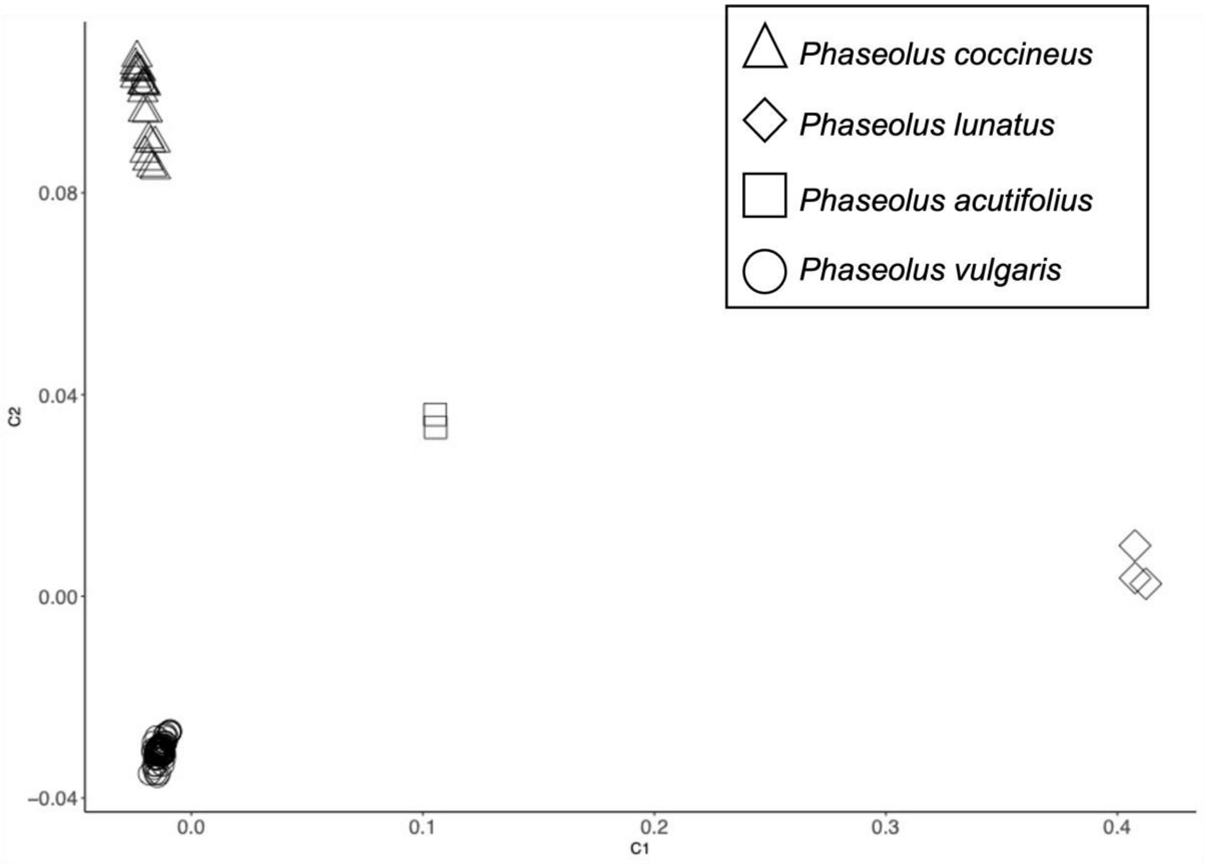

Supplementary_Figure_5: Results of the Multidimensional Scaling Analysis (MDS) of all accessions included in the analyses. Component C1 and C2 are reported in x and y axes.

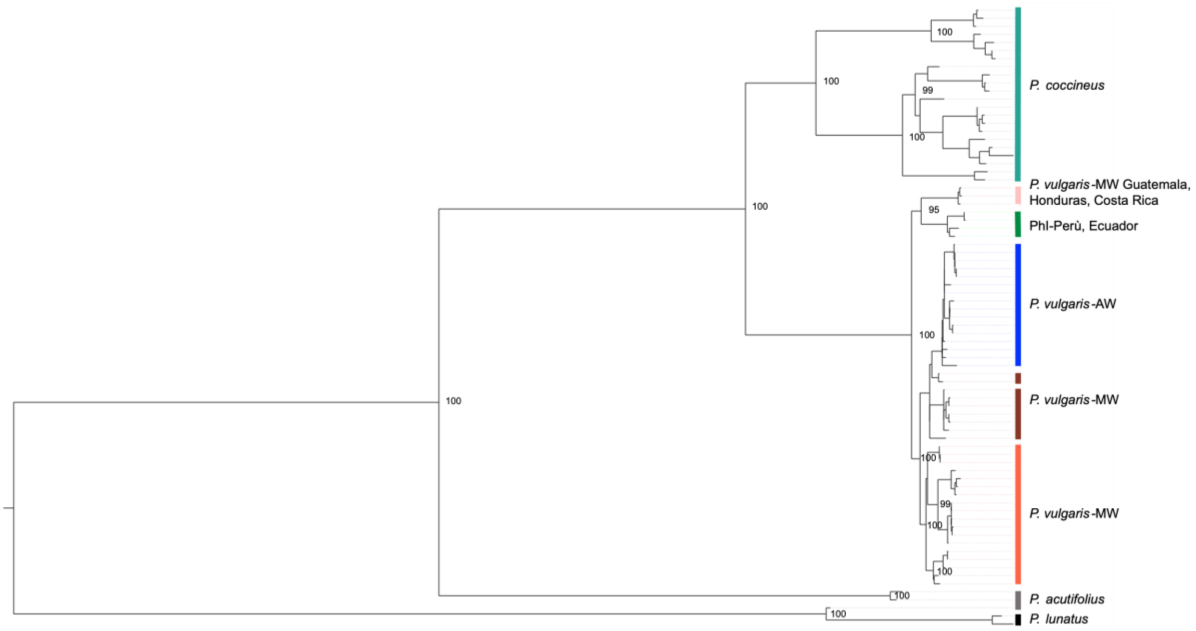


Supplementary_Figure_6: ML tree obtained from 3,231 SNPs collected from 97 samples of *Phaseolus* spp, bootstrap value=10,000.

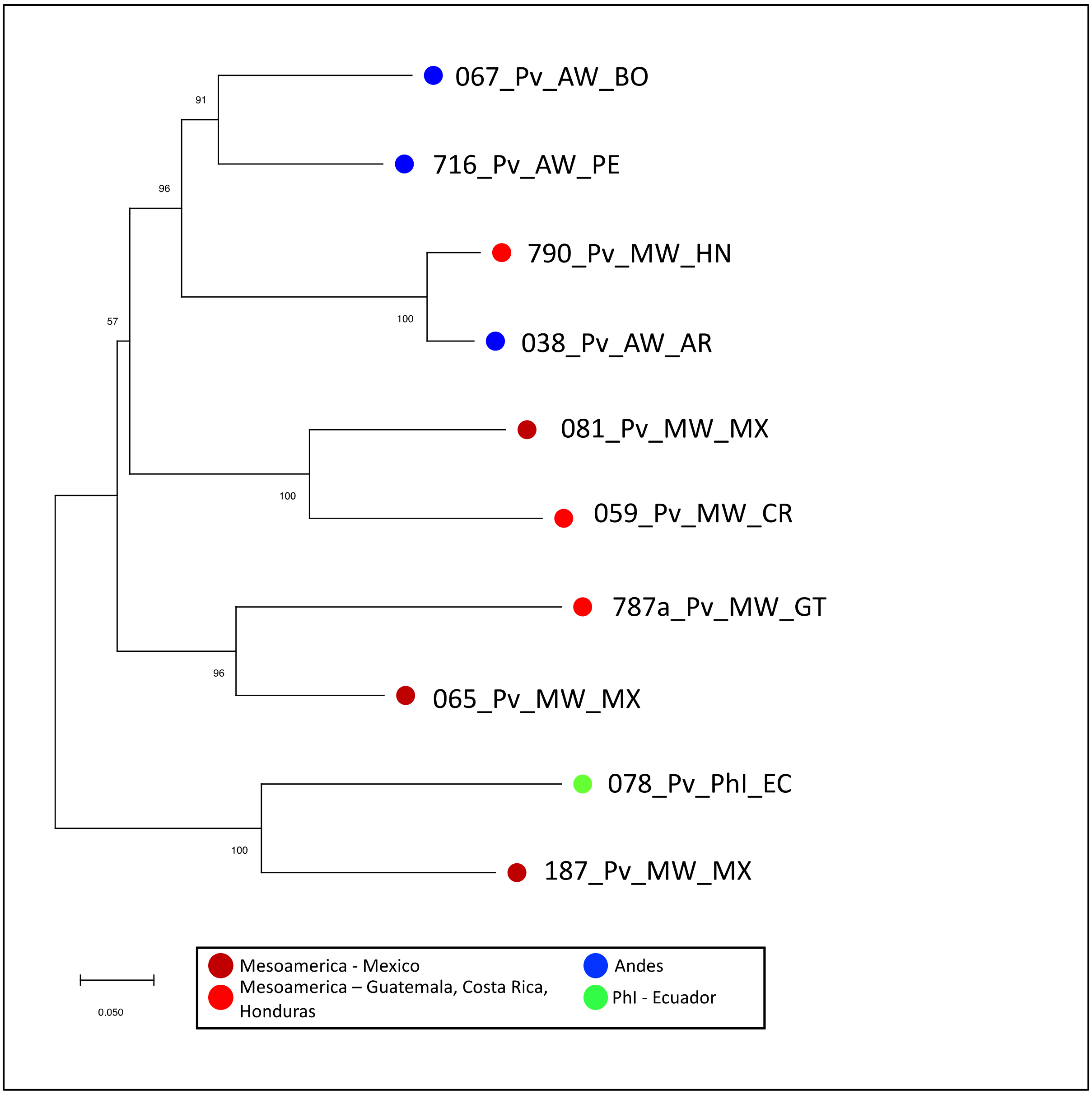


Supplementary_Figure_7: Neighbor Joining tree performed with nuclear SNPs selected every 250 kb with a bootstrap value of 10,000. In dark red: Mesoamerican samples from Mexico, light red: Mesoamerican samples from Guatemala, Costa Rica or Honduras, blue: Andean samples and in green: PhI sample from Ecuador.

### Isolation and enrichment of phaseolin and SDS-PAGE analysis

To verify the presence of the phaseolin protein type I, 7 accessions were used, 5 of which are included in the present study (i.e., G11056, G19891, G19898, G23429, G23726) and the remaining two (i.e., G21245 and G21245A) were chosen from the CIAT database due to the documented presence of the phaseoline type I.

Seed flours were extracted with 20 volumes of protein extraction buffer (Sodium Borate buffer 180 mM pH9, NaCl 0.5 M) for 2h at room temperature and then were centrifuged at 20.000 g for 20 minutes at room temperature. The supernatant fraction was acidified with the addition of 1/80 volume of glacial acetic acid and kept at 4°C for 15 minutes. This caused immediate precipitation of phaseolin. The suspension was centrifuged at 20.000 g for 20 minutes at 4°C. The precipitate was directly dissolved in denaturation buffer (20 mM TRIS-HCl pH 8.6, 1% SDS, 0.6% □ME, 0.8% Glycerol), heat denatured at 100°C for 5 min and loaded on 15% SDS-PAGE as previously described by Bollini and Chrispeels (1978) Characterization and subcellular localization of vicilin and phytohemagglutinin, the two major reserve proteins of *Phaseolus vulgaris L.* [Planta 142:291-298].

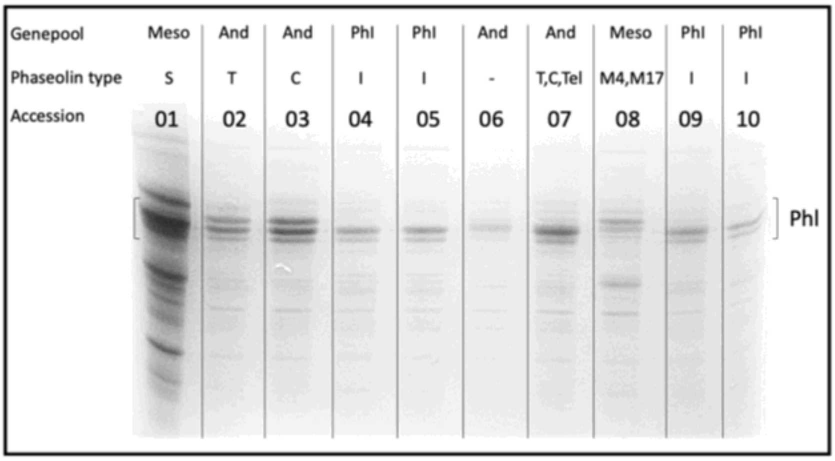


Supplementary_Figure_8: SDS-PAGE results. Accessions are listed as follow: 01, G11056; 02, G19891; 03, G19898; 04, G21245; 05, G21245; 06, G21245; 07, G21245A; 08, G23429; 09, G23726; 10, G23726.

Supplementary_Table_1: Panel of accessions selected to study the cpDNA.

| Project code | Species | Accession Number | Country |
|---|---|---|---|
| P043_Pv_MW_GT | *P. vulgaris* | G19909 | Guatemala |
| 044_Pv_MW_MX | *P. vulgaris* | G20515 | Mexico |
| P047_Pv_MW_CO | *P. vulgaris* | G21117 | Colombia |
| 069_Pv_MW_CO | *P. vulgaris* | G23462 | Colombia |
| 501_Pv_MW_MX | *P. vulgaris* | PI417671 | Mexico |
| 059_Pv_MW_CR | *P. vulgaris* | G23418 | Costa Rica |
| 746a_Pv_MW_MX | *P. vulgaris* | G12879 | Mexico |
| 787a_Pv_MW_GT | *P. vulgaris* | G23439 | Guatemala |
| 790_Pv_MW_HN | *P. vulgaris* | G50724 | Honduras |
| 887_Pv_MW_MX | *P. vulgaris* | NI1433 | Mexico |
| 911_Pv_MW_MX | *P. vulgaris* | 86 | Mexico |
| 951_Pv_MW_MX | *P. vulgaris* | M31 | Mexico |
| 716_Pv_AW_PE | *P. vulgaris* | G23458 | Peru |
| P832_Pv_MW_MX | *P. vulgaris* | G23470 | Mexico |
| 835_Pv_MW_MX | *P. vulgaris* | G23551 | Mexico |
| id837fa1_Pv_MW_MX | *P. vulgaris* | G2771 | Mexico |
| id837fa2_Pv_MW_MX | *P. vulgaris* | G11056 | Mexico |
| id837fa3_Pv_MW_MX | *P. vulgaris* | G12957 | Mexico |
| id837fa4_Pv_MW_MX | *P. vulgaris* | G13021 | Mexico |
| id837fa7_Pv_MW_MX | *P. vulgaris* | G50899 | Mexico |
| id837fa8_Pv_MW_MX | *P. vulgaris* | G12873 | Mexico |
| 007_Pv_MW_MX | *P. vulgaris* | G9989 | Mexico |
| 010_Pv_MW_MX | *P. vulgaris* | G11050 | Mexico |
| 011_Pv_MW_MX | *P. vulgaris* | G11051 | Mexico |
| 015_Pv_MW_MX | *P. vulgaris* | G12872 | Mexico |
| 016_Pv_MW_MX | *P. vulgaris* | G12877 | Mexico |
| 017_Pv_MW_MX | *P. vulgaris* | G12896 | Mexico |
| 019_Pv_MW_MX | *P. vulgaris* | G12922 | Mexico |
| 020_Pv_MW_MX | *P. vulgaris* | G12924 | Mexico |
| 021_Pv_MW_MX | *P. vulgaris* | G12927 | Mexico |
| 022_Pv_MW_MX | *P. vulgaris* | G12930 | Mexico |
| 028_Pv_MW_MX | *P. vulgaris* | G13505 | Mexico |
| 056_Pv_MW_CO | *P. vulgaris* | G22304 | Colombia |
| 057_Pv_MW_MX | *P. vulgaris* | G22837 | Mexico |
| 065_Pv_MW_MX | *P. vulgaris* | G23429 | Mexico |
| 076_Pv_MW_MX | *P. vulgaris* | G23652 | Mexico |
| 080_Pv_MW_MX | *P. vulgaris* | G24378 | Mexico |
| 081_Pv_MW_MX | *P. vulgaris* | G24571 | Mexico |

| | | | |
|---|---|---|---|
| 179_Pv_MW_MX | *P. vulgaris* | PI318696 | Mexico |
| 187_Pv_MW_MX | *P. vulgaris* | PI325677 | Mexico |
| 205_Pv_MW_MX | *P. vulgaris* | PI417775 | Mexico |
| 504_Pv_MW_MX | *P. vulgaris* | PI535430 | Mexico |
| 505_Pv_MW_MX | *P. vulgaris* | PI535409 | Mexico |
| 506_Pv_MW_MX | *P. vulgaris* | PI535450 | Mexico |
| 006_Pv_AW_AR | *P. vulgaris* | G7469 | Argentina |
| 031_Pv_AW_AR | *P. vulgaris* | G19888 | Argentina |
| 033_Pv_AW_AR | *P. vulgaris* | G19891 | Argentina |
| 038_Pv_AW_AR | *P. vulgaris* | G19897 | Argentina |
| 039_Pv_AW_AR | *P. vulgaris* | G19898 | Argentina |
| 052_Pv_AW_AR | *P. vulgaris* | G21199 | Argentina |
| 062_Pv_AW_PE | *P. vulgaris* | G23422 | Peru |
| P064_Pv_AW_PE | *P. vulgaris* | G23426 | Peru |
| 066_Pv_AW_BO | *P. vulgaris* | G23444 | Bolivia |
| 067_Pv_AW_BO | *P. vulgaris* | G23445 | Bolivia |
| 068_Pv_AW_PE | *P. vulgaris* | G23455 | Peru |
| 232_Pv_AW_AR | *P. vulgaris* | W617499 | Argentina |
| 243_Pv_AW_BO | *P. vulgaris* | W618826 | Bolivia |
| 656_Pv_AW_AR | *P. vulgaris* | G19902 | Argentina |
| 665_Pv_AW_AR | *P. vulgaris* | NI1423 | Argentina |
| 668a_Pv_AW_BO | *P. vulgaris* | G23442 | Bolivia |
| 717_Pv_AW_PE | *P. vulgaris* | G23459 | Peru |
| P718a_Pv_AW_PE | *P. vulgaris* | G23419 | Peru |
| 034_Pv_AW_AR | *P. vulgaris* | G19892 | Argentina |
| 040_Pv_AW_AR | *P. vulgaris* | G19901 | Argentina |
| 715_Pv_AW_PE | *P. vulgaris* | G23456A | Peru |
| 013_Pv_AW_PE | *P. vulgaris* | G12856 | Perù |
| id837fa6_Pv_PhI_EC | *P. vulgaris* | G23724 | Ecuador |
| 073_Pv_PhI_EC | *P. vulgaris* | G23582 | Ecuador |
| 075_Pv_PhI_PE | *P. vulgaris* | G23587 | Perù |
| 078_Pv_PhI_EC | *P. vulgaris* | G23726 | Ecuador |
| Pc_1_W_MX | *P. coccineus* | PI430189 | Messico |
| Pc_3_W_MX | *P. coccineus* | NI677 | Mexico |
| Pc_4_W_MX | *P. coccineus* | NI819 | Mexico |
| Pc_18_W_MX | *P. coccineus* | NI1120 | Mexico |
| Pc_20_W_MX | *P. coccineus* | PI325598 | Mexico |
| Pc_8_W_MX | *P. coccineus* | NI1092 | Mexico |
| Pc_21_W_MX | *P. coccineus* | NI1117 | Mexico |
| Pc_2_W_MX | *P. coccineus* | PI430183 | Messico |
| Pc_5_W_MX | *P. coccineus* | NI1265 | Mexico |

| | | | |
|---|---|---|---|
| Pc_6_W_MX | *P. coccineus* | PI417607 | Mexico |
| Pc_7_W_MX | *P. coccineus* | PI417608 | Mexico |
| Pc_9_W_MX | *P. coccineus* | PI417593 | Messico |
| Pc_10_W_MX | *P. coccineus* | PI430178 | Messico |
| Pc_11_W_MX | *P. coccineus* | PI346950 | Messico |
| Pc_12_W_MX | *P. coccineus* | NI818 | Mexico |
| Pc_13_W_MX | *P. coccineus* | NI1213 | Mexico |
| Pc_14_W_MX | *P. coccineus* | NI726 | Mexico |
| Pc_15_W_MX | *P. coccineus* | NI813 | Mexico |
| Pc_16_W_MX | *P. coccineus* | NI1122 | Mexico |
| Pc_17_W_MX | *P. coccineus* | NI1028 | Mexico |
| Pc_19_W_MX | *P. coccineus* | NI1125 | Mexico |
| Pc_22_W_MX | *P. coccineus* | NI1325 | Mexico |
| Pa_D_SV | *P. acutifolius* | PI200902 | El Salvador |
| Pa_W_MX | *P. acutifolius var. acutifolius* | PI319445 | Mexico |
| Pl_D_MX | *P. lunatus* | PI313212 | Mexico |
| Pl_W_GT | *P. lunatus* | NI1689 | Guatemala |
| Pl_W_PE | *P. lunatus* | NI1771 | Peru |

Supplementary_Table_2: Panel of accessions selected to study the nuclear DNA.

| **Project code** | **Species** | **Gene-pool** | **Country** |
|---|---|---|---|
| 187_Pv_MW_MX | *P. vulgaris* | Mesoamerican Wild | Mexico (Morelos) |
| 065_Pv_MW_MX | *P. vulgaris* | Mesoamerican Wild | Mexico (Puebla) |
| 081_Pv_MW_MX | *P. vulgaris* | Mesoamerican Wild | Mexico (Oaxaca) |
| 787A_Pv_MW_GT | *P. vulgaris* | Mesoamerican Wild | Guatemala (Santa Rosa) |
| 790_Pv_MW_HN | *P. vulgaris* | Mesoamerican Wild | Honduras (El Paraiso) |
| 59_Pv_MW_CR | *P. vulgaris* | Mesoamerican Wild | Costa Rica (San Jose) |
| 038_Pv_AW_AR | *P. vulgaris* | Andean Wild | Argentina (Tucuman) |
| 067_Pv_AW_BO | *P. vulgaris* | Andean Wild | Bolivia (Tarija) |
| 716_Pv_AW_PE | *P. vulgaris* | Andean Wild | Peru (Cuzco) |
| 078_Pv_PhI_EC | *P. vulgaris* | North Peru-Ecuador (PhI) | Ecuador (Chimborazo) |

Supplementary_Table_3: Centromeric regions and corresponding number of analyzed SNPs.

| Chr | Centromeres Dimensions (Mb) | N. SNPs |
|---|---|---|
| 1 | 7.7 | 170,779 |
| 2 | 4.6 | 113,647 |
| 3 | 2.1 | 46,248 |
| 4 | 6.5 | 163,594 |
| 5 | 7.5 | 173,932 |
| 6 | 0.1 | 2,803 |
| 7 | 13.6 | 334,210 |
| 8 | 13.9 | 294,893 |
| 9 | 4.3 | 109,616 |
| 10 | 0.7 | 16,317 |
| 11 | 1.0 | 24,544 |

### References


Bollini R., and M. J. Chrispeels. "Characterization and subcellular localization of vicilin and phytohemagglutinin, the two major reserve proteins of Phaseolus vulgaris L." Planta 142.3 (1978): 291-298.

Darling A. C., B. Mau, F. R. Blattner, N. T. Perna. Mauve: multiple alignment of conserved genomic sequence with rearrangements. Genome Res. 2004 Jul;14(7):1394-403. doi: 10.1101/gr.2289704. PMID: 15231754; PMCID: PMC442156.

Frazer K. A., L. Pachter, A. Poliakov, E. M. Rubin, I. Dubchak. VISTA: computational tools for comparative genomics. Nucleic Acids Res. 2004 Jul 1;32(Web Server issue):W273-9

Katoh K., J. Rozewicki, K. D. Yamada. MAFFT online service: multiple sequence alignment, interactive sequence choice and visualization, Briefings in Bioinformatics, Volume 20, Issue 4, July 2019, Pages 1160–1166, https://doi.org/10.1093/bib/bbx108

Rozas J., A. Ferrer-Mata, J. C. Sánchez-DelBarrio, S. Guirao-Rico, P. Librado, S. E. Ramos-Onsins, A. Sánchez-Gracia (2017). DnaSP 6: DNA Sequence Polymorphism Analysis of Large Datasets. Mol. Biol. Evol. 34: 3299-3302. DOI: 10.1093/molbev/msx248
